# Subcellular Spatial Profiling Unveils Links Between Neuritic Plaques with Selective Tau Vulnerability in Alzheimer’s Disease

**DOI:** 10.64898/2026.11.05.764692

**Authors:** Huan Li, Elaina R. Gross, Ava Devine, Akhila Gundavelli, Sean A. Prell, Maya E. Woodbury, Lin Gen, Astrid Doering, Aaron Held, Bradley T. Hyman, Alberto Serrano-Pozo, Theodore Zwang, Rachel E. Bennett, Sudeshna Das

## Abstract

Alzheimer’s disease (AD) features widespread extracellular amyloid-β plaques but selective intraneuronal tau accumulation in cortical layers II/III and V, while layer IV remains relatively resistant. Here, we combined subcellular spatial transcriptomics with co-registered Aβ and pTau immunohistochemistry to determine why neurons exposed to comparable plaque-associated pathology differ in somatic pTau accumulation. Plaque-associated neuritic stress occurred similarly across layers, but layer IV neurons showed attenuated pTau accumulation and reduced somatic expression of a shared neurite–soma stress program involving *DCTN1*, *CALM3*, *NORAD*, and *PIN1*. Trajectory analysis within layers identified tau-vulnerable and resistant neuronal states implicating calcium signaling, retrograde transport, and lysosomal function. Notably, astrocytic *APOE*–neuronal *LRP1* co-expression increased significantly along the pTau-vulnerable trajectory, and *APOE* ε4 was associated with stronger coupling of local plaque pathology to pTau-associated molecular programs. These findings reveal neuronal programs and neuron-glia interactions underlying selective tau vulnerability and resistance in AD, highlighting potential tau-targeting therapeutics.

## INTRODUCTION

Alzheimer’s disease (AD) is characterized by the accumulation of extracellular amyloid-β (Aβ) plaques, intraneuronal tau aggregates, synapse loss, and progressive neuronal degeneration. Neurofibrillary tangles (NFT), a hallmark form of tau pathology, are the strongest neuropathological correlate of neuronal loss and cognitive decline^1^. Anti-amyloid therapies slow but do not halt cognitive decline in Alzheimer’s disease (AD)^2,3^, highlighting the need for complementary strategies targeting tau pathology. Yet the mechanisms linking extracellular plaque-associated stress to intraneuronal tau accumulation remain unclear. The amyloid cascade hypothesis places Aβ upstream of tau pathology^4^, however, this model does not by itself explain the spatial and cellular selectivity of tau accumulation: Aβ plaques can be broadly distributed across the neocortex^5,6^, whereas tau accumulation and neuronal loss emerge with pronounced laminar and neuron-type specificity^7,8^. This dissociation suggests that selective tau vulnerability is not simply determined by Aβ pathology itself, but also by how distinct neuronal and glial populations respond to comparable plaque-associated stress.

Understanding this differential response requires defining the local pathological exposure through which plaques may influence neuronal tau. Emerging evidence supports a more nuanced framework in which plaques function as localized microenvironments that facilitate tau aggregation. In particular, neuritic plaques, defined by a fibrillar Aβ core encircled by a halo of dystrophic axons and dendrites, have emerged as critical sites of early tau accumulation in the neocortex^9^. These dystrophic processes may represent an early interface between extracellular Aβ pathology and intraneuronal tau, where tau becomes mislocalized, hyperphosphorylated, and prone to aggregation. In mouse models with robust amyloid deposition but limited tangle formation, phosphorylated tau species (including pTau217, pTau231, and AT8) accumulate as punctate peri-plaque structures that associate with synaptic markers (e.g., SYP, PSD95) and tau kinase activation, indicating a synaptic and neuritic origin of tau dysregulation^10^. Notably, peri-plaque synaptic accumulation of phosphorylated tau species, including p-Tau217, has been observed in the human neocortex prior to the emergence of overt NFT pathology^11^.

In parallel, experimental studies demonstrate that Aβ plaques facilitate tau-seeded aggregation by promoting local amplification of tau seeds, supporting a model in which the plaque halo acts as a permissive microenvironment for tau accumulation and amplification^12^. Consistent with this model, plaque-associated toxicity increases over the clinical course of AD^13^, whereas resilience has been linked to reduced synaptic tau hyperphosphorylation in plaque-adjacent compartments^14^. Collectively, these findings position the plaque-associated neuritic compartment as an early, dynamic, and mechanistically active site of tau pathology, preceding and potentially driving subsequent somatic tau accumulation.

A critical unresolved question is why comparable plaque-associated neuritic pathology leads to different levels of somatic tau accumulation across neurons. Plaque-associated tau may initially represent a local, compartment-restricted response within damaged axons and dendrites, whereas somatic tau burden marks a more advanced cell-intrinsic transition of tau pathology into the neuronal soma. Thus, the key question is not simply whether plaques induce tau pathology, but which local neuronal and glial states determine whether neuritic tau pathology progresses to somatic tau accumulation. This distinction is especially relevant to selective neuronal vulnerability in AD. In the neocortex, excitatory projection neurons in layers II/III (L2/3) and V (L5) are among the populations most vulnerable to tau accumulation and degeneration^15–19^, whereas other excitatory populations, including layer IV (L4) excitatory neurons, often show relative resistance despite residing within the same diseased microenvironment^20^. We therefore hypothesized that plaques generate a broadly shared neuritic stress program, but that its coupling to somatic tau accumulation is shaped by neuron-intrinsic programs and the local glial niche.

Testing this hypothesis requires resolving these molecular and cellular interactions within their spatial and pathological context. Conventional bulk and single-nucleus transcriptomic methods have provided important insights into AD-associated cell states^17–19,21^, but they dissociate cells from their anatomical position, including proximity to neuritic plaques, limiting their ability to resolve the spatial and compartment-specific processes that link extracellular plaques to intraneuronal somatic tau pathology. Subcellular spatial transcriptomics with co-registered immunohistochemistry overcomes these limitations by linking neuronal molecular states to cortical layer, distance to plaque and neighboring glial cells, and somatic tau burden within intact human tissue.

Leveraging this approach, we combined subcellular-resolution spatial transcriptomics with co-registered Aβ and AT8 staining in inferior temporal gyrus (ITG) samples from *APOE* ε3/ε3 control and *APOE* ε3/ε3 and ε4/ε4 AD donors, together with matched single-nucleus RNA-sequencing^18,22,23^. This framework enabled us to define neuronal and glial states in their native spatial and pathological context and dissect how plaque-associated neuritic stress is coupled to somatic tau accumulation in nearby neurons across vulnerable and resistant populations. We further identified neuronal and glial programs associated with tau vulnerability and resistance, functionally interrogated candidate mechanisms via tau-seeding assays in human iPSC-derived neurons, and examined how these relationships are modified by *APOE* ε4.

## RESULTS

### Spatial atlas reveals layer-specific neuronal vulnerability to tau pathology despite comparable plaque burden in human ITG

To establish a spatial atlas of cell types, their gene expression states, and their organization across cortical layers and pathological niches, we profiled inferior temporal gyrus (ITG) sections from 12 donors, including 4 control *APOE* ε3/ε3, 4 AD *APOE* ε3/ε3, and 4 AD *APOE* ε4/ε4 donors (Supplementary Table S1). We performed 10x Genomics Xenium subcellular spatial transcriptomics (5,100-gene panel; Supplementary Table S1) with co-registered immunostaining for amyloid-β (Aβ) plaques and AT8-labeled phosphorylated tau (pTau) on the same tissue sections. Matched single-nucleus RNA-sequencing (snRNA-seq) references were used for cell-type annotation^18,22,23^ (Fig. 1a). This integrated framework enabled molecular and pathological features to be resolved across cortical layers, cellular compartments, and local plaque niches.

**Figure 1.**
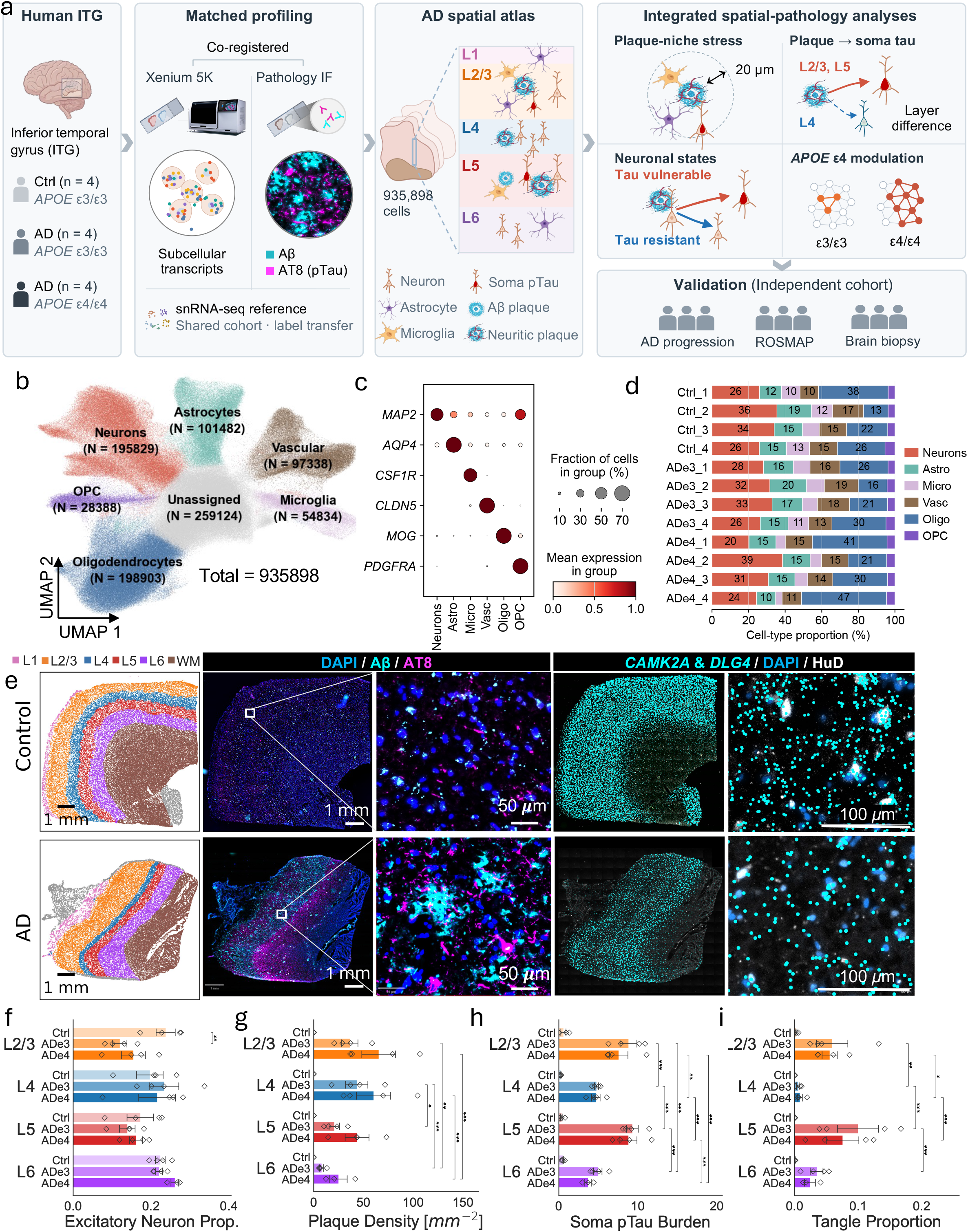
Integrated spatial transcriptomic and neuropathological profiling of human Alzheimer’s disease cortex. **a**, Experimental overview of this study. **b**, UMAP visualization showing clustering of brain cell compartments. OPC, oligodendrocyte precursor cells. **c**, Dot plot showing expression of canonical cell-type marker genes across major cell-types. Astro: astrocytes; Micro: microglia; Vasc: vascular cells; Oligo: oligodendrocytes; OPC: oligodendrocyte precursor cells. Dot size indicates the fraction of cells expressing each marker gene, and color intensity indicates the mean expression level within each cell-type. **d**, Cell-type composition across control and AD donors. Stacked bar plots show the proportion of major cell classes within each donor. **e**, Representative spatial and immunofluorescence images of control and AD inferior temporal gyrus (ITG) tissue. Left, spatial maps showing cortical layer annotations from L1 to L6 and white matter (WM). Middle panels, Aβ and AT8 co-staining showing minimal pathology in control tissue and abundant Aβ and AT8 signals in AD tissue. Right panels, Xenium transcript maps of the neuropil-enriched genes *CAMK2A* and *DLG4*, together with DAPI and HuD staining, showing abundant extrasomatic dendritic and synaptic transcripts in control cortex and marked transcript depletion in AD tissue. Scale bars, 1 mm for whole-tissue views and 50 or 100 μm for high-magnification views. **f**, Proportion of excitatory neurons in AD *APOE* ε3/ε3 and AD *APOE* ε4/ε4 vs. Ctrl and across cortical layers. **g**, Laminar distribution of Aβ plaque density, quantified as the number of plaques per mm². **h**, Soma pTau burden across cortical layers, quantified from AT8 signal within neuronal somata. **i**, Proportion of tangle-positive excitatory neurons across cortical layers. For **f**–**i**, light and dark bars indicate control and AD groups, respectively. Bars indicate the mean ± s.e.m.; Points represent individual donors (n = 4 per group). Data were analyzed with a linear mixed model (value ∼ Group × Layer) with donor as a random intercept. Pairwise comparisons were estimated as linear combinations of the model‘s fixed effects and tested with two-sided Wald z-tests. P values were adjusted for multiple comparisons with the Holm–Bonferroni procedure within each comparison family. \**q* < 0.05, \*\**q* < 0.01, \*\*\**q* < 0.001.

Across the 12 profiled brain tissue sections, 935,898 cells were captured. Unsupervised clustering of transcriptomes followed by marker-gene enrichment resolved the major cell types of the human ITG, including 195,829 neurons, 101,482 astrocytes, 54,834 microglia, 198,903 oligodendrocytes, 28,388 oligodendrocyte precursor cells (OPC), and 97,338 vascular-associated cells (Fig. 1b; Extended Data Fig. 1a). These cell-type assignments were also supported by canonical marker expression, including *MAP2* for neurons, *AQP4* for astrocytes, *CSF1R* for microglia, *MOG* for oligodendrocytes, *PDGFRA* for OPC, and *CLDN5* for vascular cells (Fig. 1c). Cell type proportions were broadly represented across all donors, indicating robust recovery of both neuronal and non-neuronal compartments across control and AD tissues (Fig. 1d; Extended Data Fig. 1b). We next leveraged matched snRNA-seq data^18^ to transfer neuronal subtypes, including layer-enriched excitatory neuronal populations: L2/3 (layer II/III), L4 (layer IV), L5 (layer V), and L6 (layer VI). Their spatial distributions recapitulated expected cortical lamination patterns, demonstrating that Xenium profiling preserved biologically meaningful layer organization in postmortem human ITG (Fig. 1e; Extended Data Fig. 1c). To link the spatial transcriptomics data to AD neuropathologic changes, the same tissue sections were co-registered with Aβ and pTau AT8 immunostaining. Control sections showed minimal Aβ and AT8 signal, whereas AD sections exhibited abundant Aβ plaques and AT8-positive tau pathology in the form of plaque-associated dystrophic neurites, NFTs, and neuropil threads (Fig. 1e). pTau pathology was prominently localized to cortical layers II/III and V in AD donors (Extended Data Fig. 1d), consistent with prior neuropathological studies demonstrating selective vulnerability of pyramidal neurons in layers II/III and layer V to tau pathology^7^.

We further examined whether the subcellular resolution of Xenium enables transcript-level readouts of neurite compartments. Neuropil-enriched transcripts^24,25^, including *CAMK2A* and *DLG4*, were readily detected in control cortex but showed marked depletion in AD tissue, both globally and in magnified regions (Fig. 1e). As these transcripts are enriched in dendrites and synapses, their reduction likely reflects loss of local dendritic and synaptic structure. Importantly, the ability to resolve transcripts outside neuronal somata enabled quantification of local transcriptomic alterations within the neuropil, including within peri-plaque niches, that are not captured by conventional snRNA-seq analyses.

Finally, we enquired whether neuronal density, Aβ plaque and tau NFT pathology showed similar laminar distributions. Vulnerable L2/3 excitatory neuronal populations showed significantly decreased abundance in AD relative to control donors, whereas no differences were observed in L4, L5 or L6 excitatory neurons (Fig. 1f). Among AD donors, Aβ plaque burden was comparable in the outer cortical layers II/III vs. IV, lower in V and VI, and higher in APOE ε4/ε4 than ε3/ε3 donors (Fig. 1g). In contrast, somatic pTau burden (computed as the neuronal soma area-normalized AT8 integrated density) was enriched in L2/3 and L5 excitatory neurons, relative to L4 and L6 excitatory neurons (Fig. 1h). Consistent with somatic pTau burden, the proportion of NFT-bearing neurons also showed layer specificity, with higher proportion of NFT-bearing neurons in tau-vulnerable L2/3 and L5 excitatory neuronal populations, and significantly lower proportions in tau-resistant L4 and L6 excitatory neurons (Fig. 1i). Thus, neurons residing in cortical layers with comparable plaque burden showed markedly different somatic pTau accumulation, establishing L2/3 and L5 excitatory neurons as tau-vulnerable and L4 as relatively tau-resistant populations.

### Layer-wise transcriptomic differences in excitatory neurons precede somatic pTau accumulation

To characterize the global transcriptomic changes of excitatory neurons across disease and pathology states, we stratified them within each cortical layer into five pathology groups: i) CTRL: control neurons; ii) AD pTau⁻ (far-plaque): AD pTau⁻ neurons located far from plaques (>50 μm); iii) AD pTau⁻ (peri-plaque): pTau⁻ neurons within peri-plaque regions (≤50 μm); iv) AD pTau⁺/Tangle⁻: AD neurons with detectable somatic pTau but not yet mature NFTs; and v) AD Tangle⁺: NFT-bearing neurons, enabling us to distinguish broad AD-associated changes from those associated with plaque proximity and progressive somatic tau pathology.

Layer-wise principal component analysis (PCA) of pseudobulk transcriptomes revealed that excitatory neurons from AD brains were transcriptionally different from CTRL (Fig. 2a; Extended Data Fig.2a). Notably, far-plaque and peri-plaque AD pTau⁻ neurons occupied similar transcriptional states across layers, suggesting that broad AD-associated transcriptional remodeling is not restricted to the immediate plaque microenvironment. Consistent with this finding, analysis of plaque distance as a continuous variable identified virtually no distance-associated differentially expressed genes (Extended Data Fig. 2b). By contrast, transcriptional variation associated with neuronal somatic tau pathology was layer dependent. AD pTau⁺/Tangle⁻ neurons were heterogeneous, with greater displacement from the pTau⁻ groups in L2/3 and L5 than in L4 and L6 excitatory neurons (Fig 2a). Tangle⁺ neurons showed further divergence from both CTRL and AD pTau⁻ states (Fig. 2a; Extended Data Fig. 2a). Thus, a broad AD-associated neuronal state precedes detectable somatic pTau.

**Figure 2.**
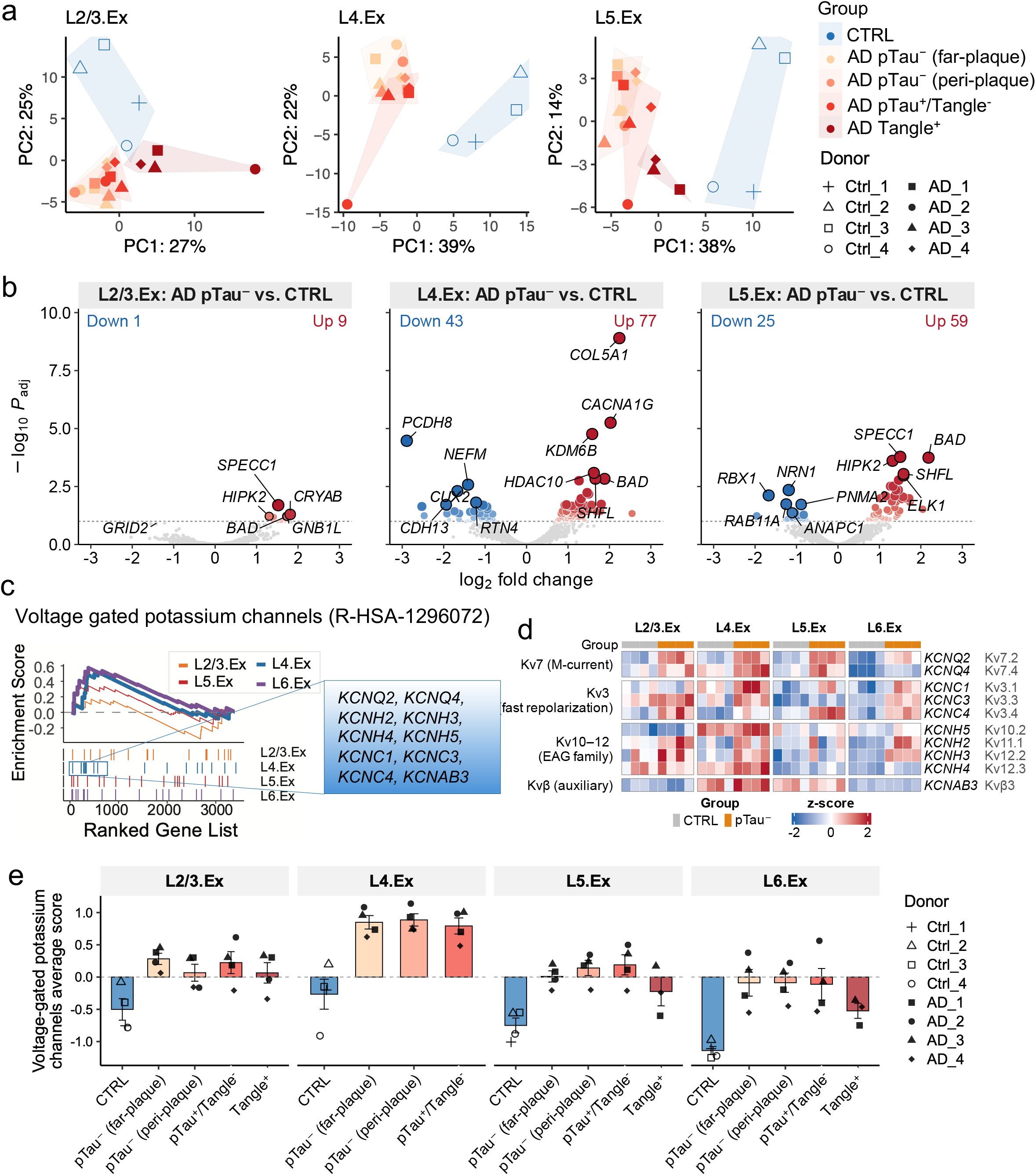
Excitatory neurons are transcriptionally dysregulated before overt pTau accumulation. **a**, Principal component analysis (PCA) of layer-wise donor-level pseudobulk transcriptomes from excitatory neurons stratified by pathology state. Within each cortical layer, neurons were grouped as CTRL (control), AD pTau⁻ (far-plaque), AD pTau⁻ (peri-plaque), AD pTau^+^/Tangle^-^, and Tangle+ (tangle-bearing) neurons. Points represent donor-level pseudobulk profiles after covariate adjustment; shaded polygons indicate groups. Each donor has a different shape. Percentages on axes indicate variance explained by each principal component. **b**, Volcano plots showing differential gene expression between AD pTau-free and CTRL excitatory neurons in L2/3, L4 and L5. Models were adjusted for batch, age and sex. Red and blue points indicate significantly upregulated and downregulated genes, respectively; selected genes are labeled. **c**, Plot showing gene-set enrichment analysis (GSEA) of ranked differential-expression signatures from AD pTau⁻ vs. CTRL. Enrichment curves are shown for voltage-gated potassium-channel gene sets across excitatory neurons. Tick marks indicate the position of genes from each set in the ranked gene list; selected leading-edge genes are highlighted. **d**, Heatmap showing the expression of selected leading-edge potassium-channel genes across CTRL and AD pTau⁻ donor-level pseudobulk samples, stratified by cortical layer. **e**, Donor-, layer-, and pathology-group-level pseudobulk expression of potassium-channel genes was normalized, gene-wise scaled across all donor–layer–group profiles, and averaged to derive the module score. Bars show the mean across donors, with individual donors overlaid as symbols.

Differential expression analysis comparing these AD pTau⁻ neurons with CTRL neurons (adjusting for age, sex, and batch) identified layer-specific transcriptional programs preceding somatic pTau accumulation (Fig. 2b; Extended Data Fig. 2c,d,e; Supplementary Table S2). Notably, pTau-resistant L4 excitatory neurons exhibited a distinct transcriptional response enriched for voltage-gated potassium-channel pathways, including *KCNQ2*, *KCNQ4*, *KCNH2/3/4/5*, *KCNC1/3/4*, and *KCNAB3* (Fig. 2c). Donor-level pseudobulk expression of the individual genes confirmed highest expression of these genes in L4 AD pTau⁻ excitatory neurons (Fig. 2d). To distinguish increased pathway expression in AD from differences in baseline expression across layers, we calculated a potassium-channel module score across donor-layer-pathology groups with pseudobulk data. L4 excitatory neurons showed increased module scores across AD pathology groups relative to CTRL, whereas the apparent enrichment in L6 excitatory neurons largely reflected its low CTRL baseline (Fig. 2e). Given the role of potassium channels in regulating membrane excitability, resting-potential stability, and spike-frequency adaptation^26,27^, this pattern suggests a layer-specific ionic response that may buffer AD-associated hyperexcitability. Thus, even before somatic pTau accumulation resistant L4 excitatory neurons engaged a distinct excitability-regulating program.

Together, these findings identify layer-specific neuronal states that precede somatic pTau accumulation and reflect subsequent tau vulnerability. The transcriptomic similarity between far-plaque and peri-plaque pTau⁻ neurons suggest that proximity to Aβ-immunoreactive deposits alone does not capture plaque-associated neurotoxicity. We therefore next examined plaque-associated neuritic pTau as a more direct measure of local plaque neurotoxicity than plaque proximity alone.

### Plaque neurotoxicity is associated with local neuritic and glial stress program across all layers

To examine plaque-associated neurotoxicity, we quantified neuritic plaque pTau burden (NP pTau) as the integrated non-somatic AT8 signal within plaques and their surrounding halo (up to 20 μm from the plaque edge), combining both signal intensity and spatial extent (see Methods). NP pTau varied independently of plaque size (Extended Data Fig. 3a, 3b), supporting its use as a measure of local plaque-associated tau pathology beyond plaque size alone. NP pTau burden was comparable across cortical layers (Fig. 3a), suggesting broadly similar plaque-associated pathological exposure across layers.

**Figure 3.**
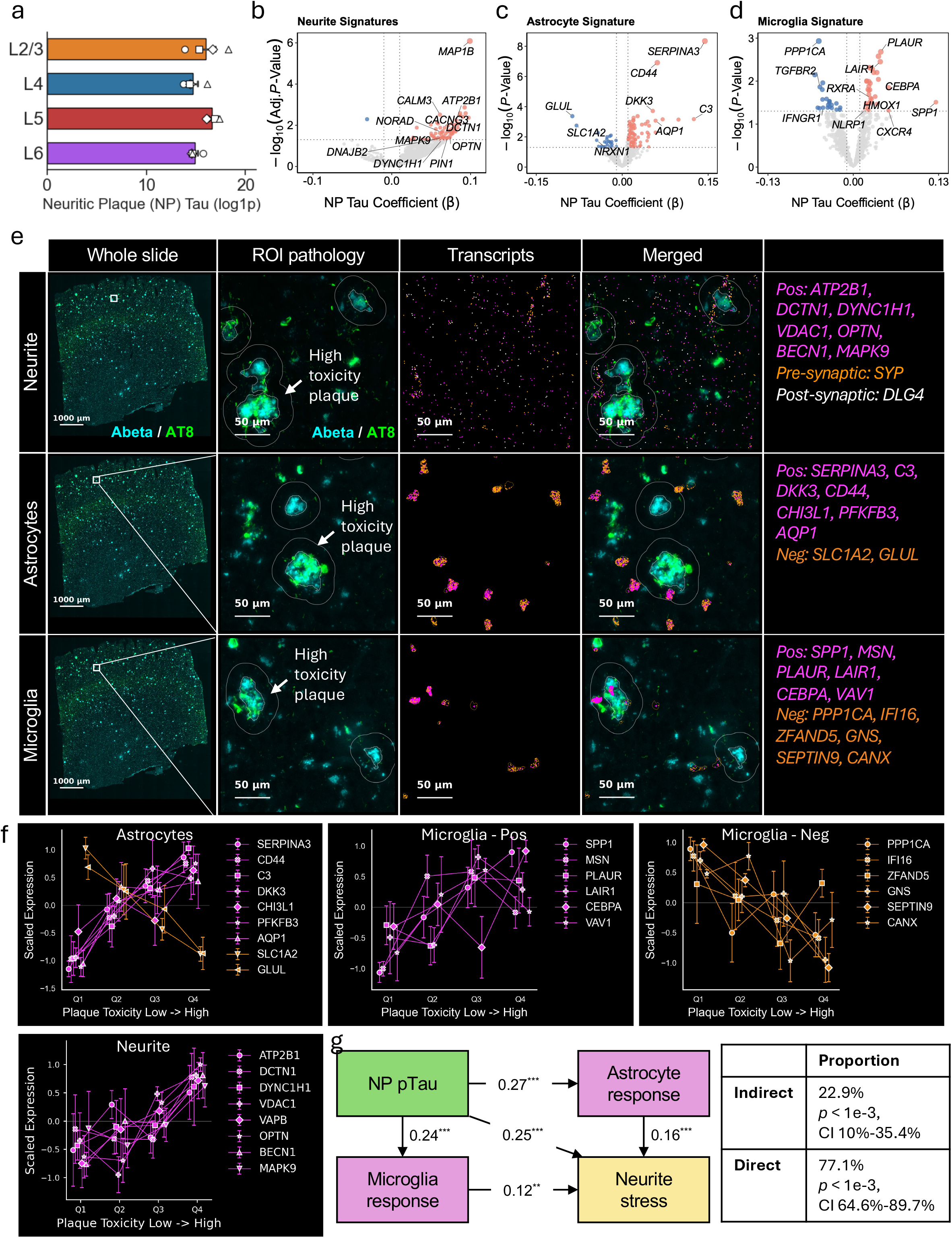
Plaque-associated toxicity drives neuritic stress predominantly through direct effects. **a**, Bar plot showing plaque-associated toxicity, measured as neuritic plaque (NP) pTau burden, across cortical layers. Bars indicate mean ± s.e.m.; each point represents the donor-level median NP pTau burden within a cortical layer, with AD donors shown using distinct symbols. **b-d**, Volcano plots showing associations between NP pTau and plaque-proximal (≤ 20 ***μ***m) transcriptional signatures in neurite (**b**), astrocytes (**c**) and microglia (**d**). The x axis shows the regression coefficient for NP pTau burden, and the y axis shows statistical significance. Red and blue points indicate positively and negatively associated genes, respectively; selected genes are labeled. **e,** Representative spatial examples showing NP pTau-associated gene responses in neuritic, astrocytic and microglial compartments. Columns, from left to right, show whole-slide Aβ/AT8 staining, example plaque-region pathology, detected transcripts, merged images and the corresponding transcript labels. Aβ is shown in cyan and AT8 in green. White outlines indicate plaque-core and halo regions. **f**, Expression of representative NP pTau-associated genes across NP pTau quartiles. Values show scaled expression for astrocytic, microglial-positive, microglial-negative and neuritic signatures. Points and error bars indicate mean ± s.e.m. across donors. Magenta and orange indicate genes positively and negatively associated with NP pTau, respectively. **g**, Structural equation model testing direct and glia-mediated effects of plaque toxicity on neuritic stress. Arrows indicate standardized path coefficients. The table summarizes the proportion of the total plaque-toxicity effect mediated indirectly through astrocytic and microglial responses versus the direct effect on neuritic stress. P values and confidence intervals are shown for indirect and direct effects.

Increasing NP pTau was associated with coordinated transcriptional alterations within the neurite compartment (Fig 3b; Supplementary Table S3). These neurite transcripts (i.e., non-somatic) within the plaque niche revealed an upregulation of gene programs consistent with neuritic stress and dysfunction (Fig 3b), including signatures of cytoskeletal and trafficking perturbation (*DCTN1, DYNC1H1, MAP1B*), calcium dysregulation and excitatory stress (*CALM3, ORAI2, ATP2B1*), synaptic impairment (*CACNG3, PTPN5*), and proteostatic imbalance (*DNAJB2, OPTN, ATP6V1H, PIN1*). Additional changes implicated mitochondrial function and metabolic stress (e.g., *VDAC1, TRAP1, DHCR24*), as well as endomembrane trafficking and organelle dynamics (e.g., *VAPB, ATL3, SNX17*), suggesting perturbations in intracellular homeostasis within plaque-proximal neuritic processes. Additionally, the upregulation of neuron-glia signaling components, such as chemokine *CX3CL1* (Supplementary Table S3), suggests potential crosstalk between neuronal processes and microglia in the plaque niche.

In parallel, NP pTau was associated with pronounced astrocytic and microglial transcriptional responses within plaque-proximal halo (20 μm) regions. Astrocytes exhibited a transcriptional shift toward a reactive state, marked by upregulation of canonical reactivity genes including *C3, SERPINA3, CHI3L1*, and *CD44*, alongside increased expression of metabolic and stress-related factors such as *PFKFB3*, *CLU*, and *MAOB* (Fig. 3c; Supplementary Table S4). This was accompanied by a coordinated downregulation of homeostatic astrocytic programs, including glutamate handling and synaptic support genes (*GLUL, SLC1A2*), as well as structural and signaling components (*SPTBN1, PTPRZ1, ADGRB1*), indicating a loss of normal astrocytic support functions (Fig. 3c; Supplementary Table S4). Microglia within the same peri-plaque regions displayed a parallel transition toward a reactive, disease-associated state, characterized by upregulation of genes involved in inflammatory signaling and cellular remodeling (e.g., *SPP1*, *CXCR4, PLAUR, VAV1, HMOX1*), coupled with downregulation of genes linked to homeostatic surveillance and intracellular trafficking (e.g., *SNX17, IFNGR1, TGFBR2, PRKACA*) (Fig. 3d, Supplementary Table S5). Representative genes in the neurite, astrocytes and microglia are shown in Fig. 3e and 3f. Together these findings show that increasing plaque toxicity is accompanied by a coordinated local response encompassing neuritic stress, glial activation, and loss of homeostatic support.

Next, to quantify direct and indirect effects of plaque toxicity on neuritic stress, we applied a structural equation modeling (SEM) framework. While plaque toxicity was significantly associated with both glial activation and neuritic stress, mediation analysis suggested that the plaque–neurite association is mostly direct (Fig. 3g). Specifically, 77.1% of the total effect was attributable to a direct glial-independent pathway between NP pTau and the neuritic stress signature (p < 1×10⁻³, 95% CI: 64.6%–89.7%), whereas a smaller fraction (22.9%) was mediated through astrocytic and microglial responses (p < 1×10⁻³, 95% CI: 10.0%–35.4%). Collectively, these findings position the plaque-associated neuritic compartment as an early, dynamic, and mechanistically active site of tau pathology, with plaque toxicity showing a strong local association with neuritic stress that was partially explained by glial responses.

### Layer IV excitatory neurons restrict somatic deployment of a plaque-associated neuritic stress program

We next asked whether local plaque toxicity translates into somatic tau accumulation and whether this relationship differs across cortical layers. For each excitatory neuron, we calculated an exponential-decay plaque toxicity exposure score integrating NP pTau from plaques within 100 μm, weighted by their distance to the neuronal soma (Fig. 4a). Plaque toxicity exposure was broadly comparable across L2/3, L4, and L5 and lowest in L6 excitatory neurons, reflecting layer VI lower plaque density and greater neuron-to-plaque distance (Fig 1g; Extended Data Fig. 4a). In contrast, somatic pTau differed markedly across layers: L2/3 and L5 excitatory neurons accumulated high pTau, whereas L4 neurons showed substantially lower pTau in their soma despite comparable plaque toxicity exposure (Fig. 4b). Consistent with this dissociation, plaque toxicity exposure was strongly associated with somatic pTau in vulnerable L2/3 and L5 excitatory neurons but significantly attenuated in L4 counterparts (Fig. 4c). Thus, comparable local plaque toxicity produces markedly different somatic pTau responses across cortical layers, identifying L4 as relatively resistant to the coupling of plaque exposure to somatic tau accumulation.

**Figure 4.**
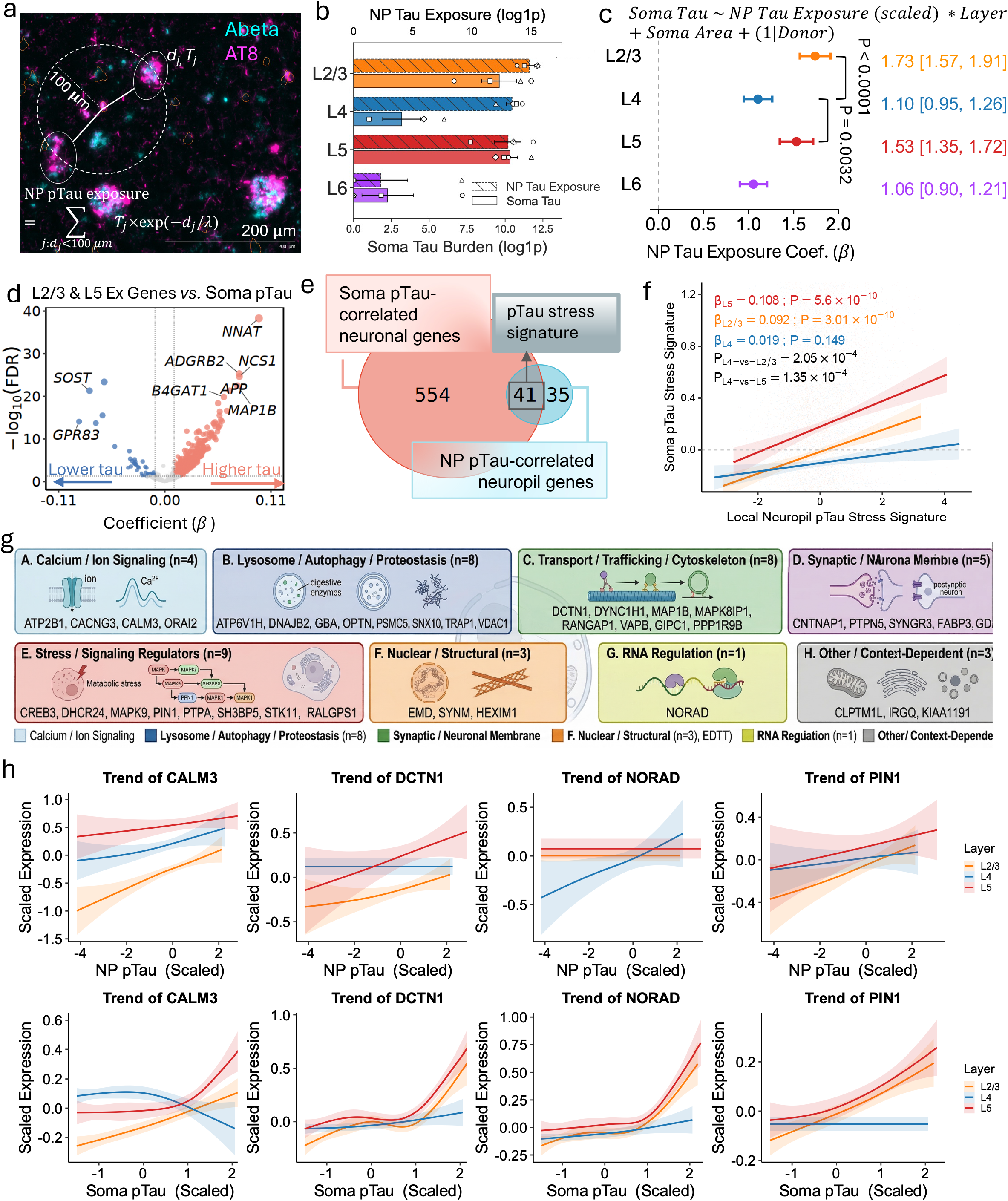
Layer 4 neurons restrict somatic deployment of a plaque-associated stress program. **a**, Schematic showing the calculation of neuron-level NP pTau exposure. **b**, Bar plot showing NP pTau exposure and somatic pTau burden across excitatory neuron layers. Filled bars show NP pTau exposure; open bars show soma pTau burden. Bars indicate mean ± s.e.m.; points represent individual donors, with AD donors shown using distinct symbols. **c**, Forest plot showing layer-specific association between NP pTau exposure and soma pTau burden. Points indicate estimated coefficients; horizontal lines indicate 95% confidence intervals. P values indicate layer comparisons. **d**, Volcano plot showing genes associated with soma pTau burden in vulnerable L2/3 and L5 excitatory neurons. The x axis shows the regression coefficient for soma pTau burden, and the y axis shows statistical significance. Red and blue points indicate genes positively and negatively associated with soma tau burden, respectively; selected genes are labeled. **e**, Overlap between soma pTau-correlated neuronal genes and NP pTau-correlated neuropil genes. The intersecting set defines a shared pTau stress signature linking plaque-associated neuritic responses with somatic tau-associated transcriptional programs. **f**, Association between neuritic and somatic pTau stress signatures across excitatory neuron layers. Lines show fitted relationships between the NP pTau-correlated neuritic signature and the soma pTau-correlated neuronal signature; shaded areas indicate 95% confidence intervals. Regression coefficients and P values are shown for each layer, together with pairwise comparisons between layers. **g**, Functional grouping of genes in the shared pTau stress signature. **h**, Representative layer-specific expression trends of shared pTau stress genes NP pTau or soma pTau in L2/3, L4 and L5 excitatory neurons. Lines indicate fitted trends; shaded areas indicate 95% confidence intervals.

To investigate molecular programs associated with this coupling, we identified genes associated with somatic pTau in vulnerable L2/3 and L5 excitatory neurons (Fig. 4d; Supplementary Table S6) and compared them with neuritic stress genes, i.e. those associated with NP pTau (Fig. 3b). Of 76 NP pTau-associated neuritic genes, 41 were also associated with somatic pTau, representing significant enrichment over chance (hypergeometric P = 4.6 × 10⁻⁴; 1.48-fold enrichment; Fig. 4e), and the two programs were significantly correlated across L2/3 and L5 excitatory neurons (Fig. 4f; Extended Data Fig. 4b). However, in L4 excitatory neurons the correlation was not statistically significant and was attenuated compared with vulnerable L2/3 and L5 excitatory neurons (Fig. 4f). The common 41 genes represented a coordinated transcriptional response encompassing calcium and ion signaling (*ATP2B1, CACNG3, CALM3, ORAI2, VDAC1*), lysosomal and proteostatic pathways (*ATP6V1H, DNAJB2, GBA, OPTN, PSMC5, SNX10, TRAP1)*, intracellular transport and cytoskeletal dynamics (*DCTN1, DYNC1H1, MAP1B, MAPK8IP1, RANGAP1, VAPB, GIPC1, PPP1R9B*), and synaptic and membrane-associated processes (*CNTNAP1, PTPN5, SYNGR3, FABP3, GDA*). Additional enrichment was observed for stress signaling regulators (*CREB3, DHCR24, MAPK9, PIN1, PTPA, SH3BP5, STK11, ZPR1, RALGPS1*), nuclear and structural components (*EMD, SYNM, HEXIM1*), and RNA regulatory elements (*NORAD*), consistent with a broad neuronal stress-response program (Fig. 4g). Together, these genes define a shared neurite-to-soma stress program associated with progression from local plaque toxicity to somatic pTau accumulation.

Representative genes from the shared neurite–soma stress program, including CALM3, DCTN1, NORAD, and PIN1, showed increased expression with NP pTau across layers, including L4 (Extended Data Fig. 4c). However, their expression increased with somatic pTau primarily in vulnerable L2/3 and L5 neurons, with attenuated responses in L4, suggesting restricted somatic deployment of the neuritic stress program (Fig. 4h). This differential coupling was present in all 41 genes (Extended Data Fig. 4d). Thus, L4 excitatory neuronal resistance is not explained by reduced plaque toxicity or absence of local neuritic stress, but is associated with restricted somatic deployment of a shared plaque-associated stress program.

### Neuronal states and neuron-glia signaling are associated with differential somatic pTau accumulation

Next, we investigated whether excitatory neurons within each layer differ in their propensity to accumulate somatic pTau at comparable levels of NP pTau exposure. Towards that end, we performed layer-specific trajectory analysis of L2/3, L4, and L5 excitatory neurons using transcriptomic profiles alone, and subsequently mapped NP pTau exposure, somatic pTau burden, and NFT-bearing status onto the inferred trajectories. Across layers, trajectories bifurcated from a common State 1 into two transcriptionally distinct branches with overlapping NP pTau exposure but divergent somatic pTau burden (Fig. 5a–d). State 3 showed progressively greater somatic pTau along pseudotime, with NFT-bearing neurons concentrated at later pseudotime (Extended Data Fig. 5a–d). In contrast, State 2 showed significantly lower somatic pTau despite overlapping NP pTau exposure (Fig. 5c, d), identifying candidate pTau-vulnerable (State 3) and pTau-resistant (State 2) trajectories.

**Figure 5.**
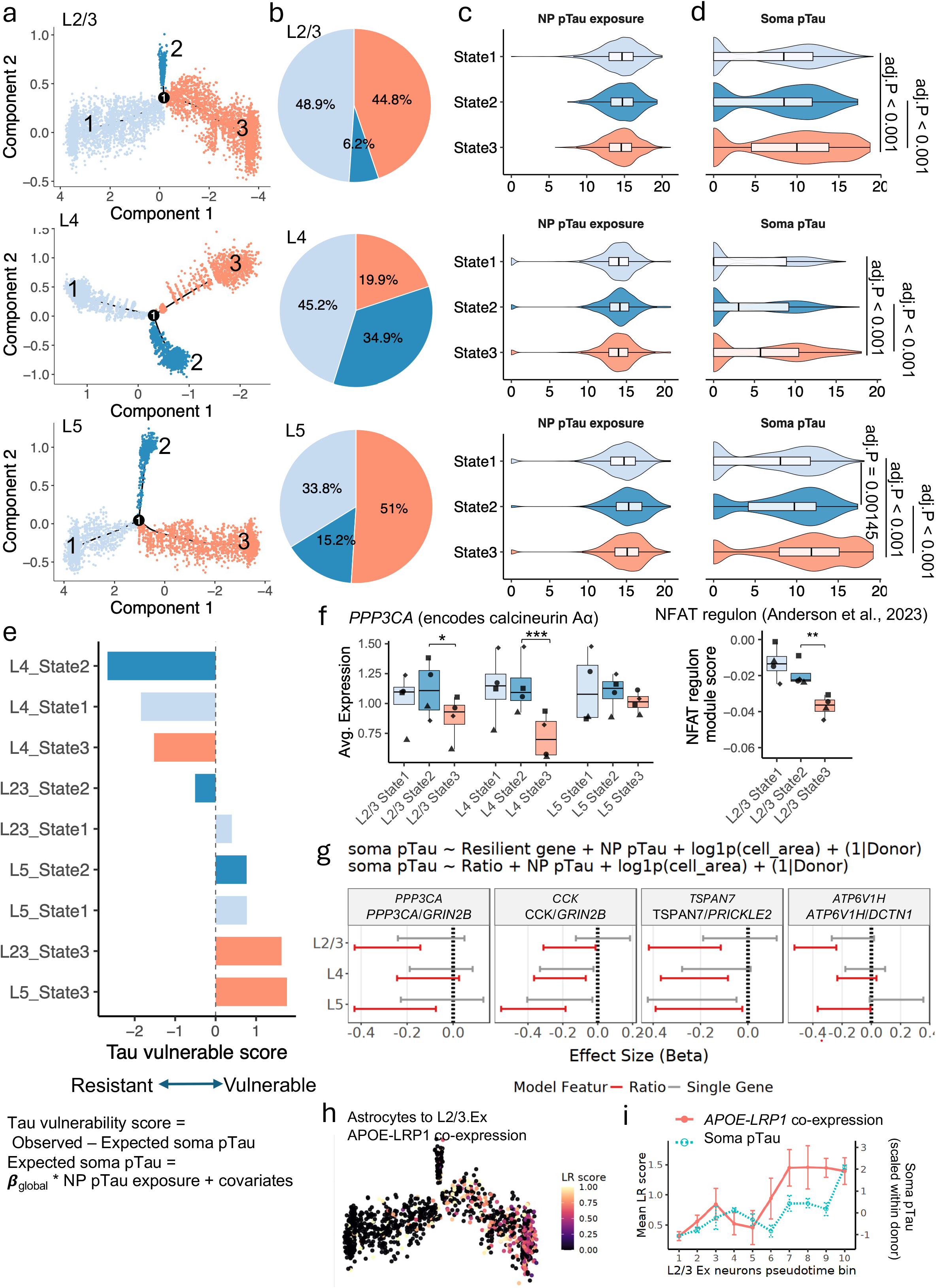
Intrinsic neuronal states modulate the conversion of plaque-associated stress into somatic pTau accumulation. **a**, Layer-specific pseudotime trajectories of L2/3, L4 and L5 excitatory neurons. Cells are colored by inferred transcriptional state. **b**, Proportion of excitatory neurons assigned to each pseudotime state in L2/3, L4 and L5. **c,d**, Distribution of NP pTau exposure (**c**) and soma pTau burden (**d**) across pseudotime states in each layer. Violin plots show cell-level distributions; embedded box plots indicate median and interquartile range. Adjusted P values are shown for state comparisons. **e**, Bar plot showing tau vulnerability score across layer-specific pseudotime states. Positive values indicate higher-than-expected soma pTau, whereas negative values indicate lower-than-expected soma pTau. **f**, Box plot showing expression of PPP3CA, which encodes the calcineurin catalytic subunit Aα (left), and activity scores for the downstream NFAT regulon derived from Anderson et al. (right), across layer–state groups. Each point represents the mean value for one donor within the neuronal population&state group. P values in the right panel were calculated using two-sided paired t-tests comparing State 2 and State 3 within each layer. *P < 0.05, **P < 0.01, ***P < 0.001, n.s., not significant. **g**, Association between soma pTau burden and resistant-to-vulnerable gene ratios across excitatory neuron layers. Forest plots show effect sizes from models testing either individual resistant genes or resistant-to-vulnerable gene ratios, adjusted for soma area and donor. Points indicate estimated coefficients; horizontal lines indicate 95% confidence intervals. Negative coefficients indicate association with lower soma pTau burden. **h**, Association of astrocyte-to-L2/3 excitatory-neuron APOE–LRP1 co-expression with somatic pTau. Left, L2/3 excitatory neurons positioned along the inferred trajectory and colored by the local APOE– LRP1 ligand–receptor co-expression score. **i**, Mean APOE–LRP1 co-expression score and somatic pTau burden across neuronal pseudotime bins; somatic pTau values were scaled within donor.

The distribution of these states mirrored laminar tau vulnerability. The pTau-resistant State 2 was most abundant in layer IV (34.9%), compared with 6.2% in layers II/III and 15.2% in layer V, whereas the pTau-vulnerable State 3 predominated in layers II/III (44.8%) and layer V (51.0%) (Fig. 5b). To quantify this relationship directly, we defined tau vulnerability as “observed minus expected” somatic pTau, with expected pTau modeled from NP pTau exposure and covariates across neurons. Positive residuals indicate greater-than-expected somatic pTau and negative residuals relative resistance. State 3 showed greater pTau vulnerability across layers, whereas State 2 showed lower vulnerability, with layer IV State 2 exhibiting the strongest resistance (Fig. 5e).

We investigated whether resistant states shared a transcriptional program across layers. Differential expression analysis comparing the pTau-resistant State 2 with the pTau-vulnerable State 3 within each layer identified a set of genes enriched in resistant states in layers II/III and V (Extended Data Fig. 5e-f; Supplementary Table S7). Genes consistently enriched in pTau-resistant State 2 included *CCK, CCT7, EEF1A2, NPTN, PCDH10, PGK1, PPP3CA, SESN3, TRRAP, TSHZ2, TSPAN7*, and *VSNL1*, implicating synaptic connectivity, calcium signaling, metabolism, and proteostasis. Conversely, pTau-vulnerable states were enriched for calcium-, stress-, and proteostasis-associated genes, including *GRIN2B*, *DHCR24*, *PIN1*, *PRICKLE2*, and *SH3BP5*, as well as the non-coding RNAs *MEG3*, *NORAD*, and *MIAT*. The vulnerable-state program showed strong concordance with the neurite-to-soma stress signature identified in Fig. 4: among the 41 genes comprising this signature, 24 were relatively higher in the vulnerable state in L2/3, 26 in L4 and 38 in L5 excitatory neurons (Supplementary Table S7).

Among resistance-associated genes, *PPP3CA*, encoding the catalytic α subunit of calcineurin, was consistently higher in State 2 than State 3 across layers (Fig. 5f). Calcineurin can dephosphorylate tau^28^ and activates NFAT signaling through dephosphorylation and nuclear translocation. We quantified NFAT regulon activity using excitatory-neuron target genes inferred from paired transcriptomic and chromatin-accessibility data reported by Anderson et al^29^. NFAT regulon activity was significantly higher in pTau-resistant State 2 in L2/3 excitatory neurons (Fig. 5f). These findings link higher *PPP3CA* expression to increased NFAT regulon activity in pTau-resistant neurons in layers II/III, with weaker or absent evidence for this relationship in other layers. Notably, ratios of resistance- to vulnerability-associated genes showed stronger negative associations with somatic pTau than individual genes, including *PPP3CA*/*GRIN2B*, *CCK/GRIN2B*, *TSPAN7/PRICKLE2*, and *ATP6V1H*/*DCTN1*, particularly in L2/3 and L5 neurons (Fig. 5g). These findings support a capacity–demand model in which tau resistance reflects the balance between resistance-associated programs and vulnerability-associated stress responses.

Although L2/3 excitatory neurons were predominantly tau-vulnerable (State 3), a 6.2% subset were classified as predominantly tau-resistant (State 2). To confirm the existence of a tau-resistance gene expression program in L2/3 excitatory neurons, we next evaluated the tau-resistance program in L2/3 excitatory neurons from independent human AD datasets: an AD progression study^18^, ROSMAP^17^, and a brain-biopsy study^16^. The L2/3 resistance module was significantly associated with greater preservation of L2/3 neurons in both the AD progression (Extended Data Fig. 5g) and ROSMAP cohorts (Extended Data Fig. 5h). Consistent with a relationship to tau resistance, the shared resistance module was lower in Aβ+/tau+ than in Aβ+/tau− donors in the brain-biopsy cohort (Extended Data Fig. 5i).

Finally, we examined whether vulnerable-state progression was coordinated with the local glial niche. Given the established role of the APOE receptor LRP1 in neuronal tau uptake and processing^30,31^, we quantified spatial co-expression between neuronal *LRP1* and *APOE* in astrocytes within 20 µm. Spatially defined *APOE*–*LRP1* co-expression was elevated toward the distal portion of the L2/3 pTau-vulnerable trajectory (Fig. 5h). Furthermore, pseudotime-binned profiles revealed a sustained increase in spatial *APOE*–*LRP1* co-expression at earlier pseudotime positions than the increase of somatic pTau (Fig. 5i). This pattern nominates *APOE–LRP1* coupling as a potential early glial correlate of neuronal tau vulnerability. Together, these findings identify transcriptionally distinct neuronal states with differential vulnerability to somatic pTau accumulation. L4 excitatory neurons are enriched in a pTau-resistant state characterized by lower-than-expected somatic pTau, whereas tau-vulnerable states are associated with neuronal stress programs and local neuron–glia signaling.

### APOE ε4 is associated with broader molecular coupling to somatic pTau accumulation

Finally, we asked whether the coupling of local plaque pathology to somatic pTau varied with *APOE* genotype. We first defined the pathways associated with NP pTau-to-somatic pTau coupling in AD neurons from *APOE* ε3/ε3 donors, focusing on the vulnerable layers II/III and V. For each gene set, we tested its association with NP pTau exposure and subsequently with somatic pTau after accounting for NP pTau, layer, and donor. Prominent soma-associated pathways included postsynaptic organization (*DLG4, SHANK3, CAMK2A, CAMK2B, GSK3B*), neuronal projection organization (*APOE, APP, BIN1, DCTN1, CDK5, GRIN2B*), calcium handling and membrane excitability (*ATP2A2, CACNA1C, CALM3, HCN4, KCNJ2, SLC8A1, TRPM4*), GnRH signaling, and regulation of peptidyl-threonine phosphorylation (*APP, GSK3A, MAPK1, RPTOR, SIRT2*) (Fig. 6a). These pathways showed stronger correlations with somatic pTau in vulnerable than resistant neuronal states.

**Figure 6.**
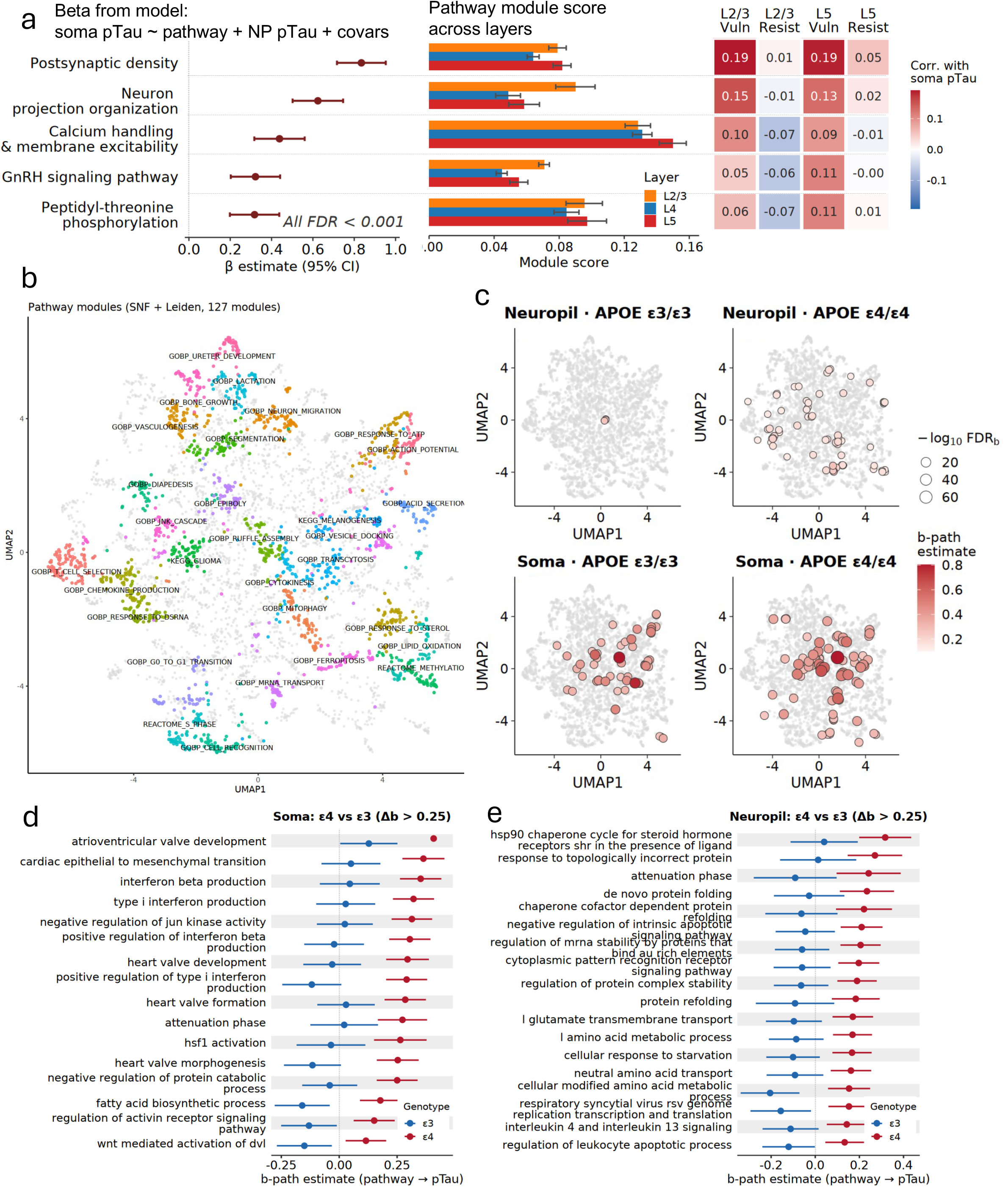
*APOE* ε4 is associated with broader pathway coupling between local plaque pathology and somatic pTau accumulation. **a,** Five soma-derived pathways associated with somatic pTau in *APOE* ε3/ε3 AD neurons from L2/3 and L5. Left, pathway coefficients from linear models relating somatic pTau to pathway score after adjustment for NP pTau exposure, layer and donor. Points denote β estimates and horizontal lines indicate 95% confidence intervals; all displayed associations had Benjamini–Hochberg false-discovery rates (FDRs) below 0.001. Middle, mean pathway scores across L2/3, L4 and L5; error bars indicate s.e.m. across donor-level averages. Right, mean within-donor Spearman correlations between pathway score and somatic pTau along the vulnerable (State 1 to State 3) and resistant (State 1 to State 2) trajectories. **b,** UMAP representation of the pathway-similarity network constructed by integrating gene-set overlap with donor- and layer-residualized pathway co-expression. Pathways were grouped into 127 modules using Leiden clustering. Colors distinguish the 30 largest modules, grey denotes the remaining modules and labels indicate representative pathway annotations. **c,** Distribution of pathways associated with plaque-to-soma pTau coupling in the local neuropil (top) and somatic compartment (bottom) of *APOE* ε3/ε3 and ε4/ε4 AD neurons. All panels use the common pathway-network embedding shown in **b**. Highlighted pathways were positively associated with both NP pTau exposure (a-path β > 0.01, FDR < 0.05) and somatic pTau after adjustment for NP pTau exposure, layer and donor (b-path β > 0.25 for somatic pathways or β > 0.10 for neuropil pathways; FDR < 0.05). Point colour represents the b-path coefficient and point size represents −log₁₀(FDR); grey points denote pathways not meeting these criteria. **d,e,** Genotype-stratified b-path coefficients for somatic (**d**) and neuropil (**e**) pathways showing an ε4/ε4–ε3/ε3 coefficient difference greater than 0.25, a positive coefficient in ε4/ε4 neurons and FDR < 0.05 in the ε4/ε4 analysis. Points denote β estimates and horizontal lines indicate 95% confidence intervals; blue and red indicate ε3/ε3 and ε4/ε4, respectively. These panels show effect-size differences between genotype-stratified models rather than formal genotype-by-pathway interaction tests. Data were obtained from four biologically independent AD donors per genotype. All reported FDRs were calculated using the Benjamini–Hochberg procedure.

We next organized pathways into a network of 127 functionally related modules based on gene overlap and gene co-expression (Fig. 6b). Mapping pTau-associated pathways onto this network revealed broader pathway engagement in *APOE* ε4/ε4 than ε3/ε3 AD neurons, particularly in the local neuropil (Fig. 6c; Supplementary Table S8). Pathways meeting both NP-pTau and somatic-pTau association criteria increased from 53 to 86 in *APOE* ε4/ε4 neuronal somata and from 3 to 70 in *APOE* ε4/ε4 neuropil, with ε4-associated pathways distributed across multiple functional modules rather than a single biological process.

Direct effect-size comparisons identified 16 somatic and 18 neuropil pathways with stronger associations with somatic pTau in ε4/ε4 vs. ε3/ε3 neurons (Δβ > 0.25; Fig. 6d,e). Neuropil differences prominently involved protein folding and chaperone activity, protein-complex stability, RNA regulation, amino-acid and glutamate transport, and cellular stress responses, whereas somatic differences included stress and proteostatic pathways together with inflammatory and lipid-related programs.

Thus, *APOE* ε4 homozygosis was associated with broader molecular coupling between local NP pTau and somatic pTau, with the strongest expansion occurring in the neuropil. These findings suggest that ε4-associated tau vulnerability reflects coordinated engagement of multiple local stress and proteostatic programs rather than a single dominant pathway.

## DISCUSSION

Using subcellular spatial transcriptomics with co-registered Aβ and AT8 immunohistochemistry, we define a multistep framework linking plaque-associated tau-immunoreactive neuritic dystrophies to somatic neuronal tau accumulation in AD. We show that i) excitatory neurons undergo transcriptional remodeling before overt somatic pTau accumulation; ii) exposure to neuritic plaques induces a coordinated local stress response in surrounding excitatory neurons; iii) the coupling of this neuritic stress to somatic pTau differs across cortical layers, with relative attenuation in layer IV (L4); and iv) this resistance neuronal state is enriched in L4; and v) *APOE* ε4 is associated with a broader set of molecular programs linking neuritic plaque neuronal exposure to somatic pTau burden. Together, these findings suggest that while plaque-associated neuritic stress is broadly shared, selective tau vulnerability reflects neuronal programs that determine whether local pathological stress progresses from the neurites to the neuronal cell body, with *APOE* ε4 amplifying this response.

Our findings indicate that the major transcriptional inflection point in excitatory neurons occurs before overt somatic pTau accumulation. In AD donors, pTau⁻ neurons were transcriptionally distinct from control neurons regardless of plaque proximity, indicating that early neuronal dysfunction extends beyond the peri-plaque niche and reflects broader disease-associated cortical remodeling. Notably, relatively pTau-resistant L4 excitatory neurons exhibited a distinct transcriptional response enriched for voltage-gated potassium channels. This finding is consistent with recent evidence that L4 excitatory neurons are relatively preserved during AD progression, with KCNIP4 implicated in regulating AD-associated neuronal hyperexcitability^20^. The enrichment of KCNQ, KCNH, and KCNC family members in our study further suggests that modulation of intrinsic neuronal excitability may contribute to L4 resistance to somatic pTau accumulation.

We identified a spatially localized neuronal response to Aβ plaque pathology within the neuritic compartment. Prior studies have identified the plaque-associated neurites as an early and mechanistically active site of pathological signaling^9–12^. We extend this framework by defining a coordinated neuritic transcriptional response to increasing plaque-associated tau burden, involving cytoskeletal and trafficking dysfunction, calcium dysregulation, synaptic impairment, and proteostatic stress. Although astrocytes and microglia mounted robust reactive responses within the same plaque-adjacent microenvironment, mediation analysis indicated that most of the plaque–neuritic stress association was direct, with glial responses accounting for a smaller indirect component. Whereas prior spatial transcriptomic studies have primarily characterized plaque-associated transcriptional changes in glia cells^32–36^, and a recent study further identified pathology-proximal transcriptional alterations in neurons^37^, our subcellular analysis directly resolves transcriptional responses within plaque-associated neuronal processes. Together, these findings suggest that neurites are primary local sufferers of plaque-associated injury, while glial activation may modulate or amplify this response.

NFT formation begins in the apical dendrites of pyramidal excitatory neurons^38,39^ and plaque-associated dystrophic neurites are sites of early tau seeding^12^ and remain continuous and connected to viable cell bodies^40^. Thus, pTau aggregation (and its downstream pathogenic cascade) could conceivably propagate from plaque-associated neuritic compartments to neuronal somata, ultimately leading to the formation of a mature NFT. We compared the pTau burden within plaque-associated neurites vs. neuronal cell bodies across layers and found that, despite broadly similar plaque-associated neuritic pathology across cortical layers, the relationship between local plaque neuritic change and somatic pTau accumulation was strongly shaped by laminar identity. Prior neuropathological and single-cell studies have established the selective vulnerability of L2/3 and L5 excitatory neurons^7,8^, and identified molecular signatures associated with neuronal resilience and vulnerability^19,20,41^. Our molecular signatures of tau vulnerability and resistance are consistent with other studies^20,42^.Our study extends this work by positioning these differences within the pathological sequence from plaque-associated neuritic stress to somatic pTau accumulation and highlighting L4 neuronal resilience to this process. Increasing plaque-associated dystrophic neurites were strongly associated with somatic neuronal pTau in vulnerable L2/3 and L5 neurons, whereas this relationship was markedly attenuated in L4 neurons. Importantly, L4 neurons still exhibited the plaque-associated neuritic stress response, indicating that resilience is not simply attributable to reduced upstream pathological exposure. Instead, L4 neurons experience and respond to plaque-associated stress but show reduced coupling of this response to somatic pTau accumulation. Thus, selective tau neuronal vulnerability reflects a combination of exposure to neuritic plaques and intrinsic neuronal mechanisms that determine whether local neuritic stress progresses to somatic tau accumulation and, ultimately, mature NFT formation^43^.

Trajectory and differential-expression analyses further identified distinct neuronal states representing divergent responses to plaque-associated neuritic stress. The resistant state, most abundant in layer IV, was enriched for programs supporting synaptic maintenance, calcium adaptation, metabolism, and proteostasis, whereas vulnerable states predominant in layers II/III and V showed greater excitatory receptor signaling, calcium-entry-associated stress, cytoskeletal remodeling, and proteostatic dysfunction. The enrichment of *PPP3CA* and NFAT activity in resistant neurons is particularly notable given the role of calcineurin in tau dephosphorylation and calcium-dependent signaling^44^. Notably, resistance was better captured by the balance between homeostatic and stress-associated programs than by individual genes alone. Ratios such as *PPP3CA*/*GRIN2B*, *CCK/GRIN2B*, *TSPAN7/PRICKLE2*, and *ATP6V1H*/*DCTN1* were associated with lower somatic pTau, suggesting that vulnerability reflects the balance between cellular buffering capacity and stress demand. Together, these findings support a capacity–demand model in which neuronal tau vulnerability depends on whether homeostatic programs are sufficient to contain plaque-associated neuritic stress and limit its progression to somatic tau pathology.

Mechanistically, several components of the shared neurite–soma program point to plausible pathways linking plaque-associated neuritic injury to somatic tau pathology. Enrichment of dynein–dynactin components, including *DCTN1* and *DYNC1H1*, implicates retrograde transport as a potential conduit linking plaque-proximal neuritic stress to more distal somatic responses. Convergence on calcium-regulatory genes further suggests that disrupted Ca²⁺ homeostasis may amplify this coupling through excitotoxic signaling, while shared induction of proteostasis regulators such as *PIN1*, *DNAJB2*, and *GBA* indicates impaired tau handling across neuritic and somatic compartments. Notably, these transport, calcium-signaling, and proteostatic programs showed stronger coupling to somatic pTau in *APOE* ε4/ε4 donors, suggesting that the ε4 allele may amplify the molecular response linking local plaque-associated neuritic stress to somatic tau accumulation. Together, these observations support a model in which neuritic stress and somatic tau accumulation are linked through coordinated transport, calcium-signaling, and proteostatic pathways, rather than representing independent compartmental responses.

A limitation of this study is that the spatial analyses are based on cross-sectional postmortem human tissue, which limits direct inference of the temporal and causal sequence linking plaque-associated neuritic stress to somatic tau accumulation. Our perturbation experiments provide functional support for one prioritized candidate gene and demonstrate compartment-specific effects on tau pathology, but do not fully recapitulate or establish causality across the plaque–neurite–soma sequence. In addition, the Xenium panel provides targeted rather than whole-transcriptome coverage, potentially limiting detection of additional pathways contributing to neuronal vulnerability or resilience. Our analysis was confined to the ITG and advanced Braak NFT stages, and it remains to be determined whether the same laminar resistance programs generalize across other neocortical regions or disease stages. Finally, while our spatial and functional findings implicate retrograde transport, calcium signaling, and proteostatic balance as candidate mechanisms, targeted perturbation of these pathways and direct measurements of tau transport, phosphorylation, and accumulation across neuronal compartments will be needed to establish how they regulate the progression from local plaque-centered neuritic stress to somatic tau pathology.

Overall, our study provides a spatially resolved framework linking plaque-associated neuritic pathology to selective somatic tau accumulation in AD. By separating pathological plaque exposure, neuritic stress, and somatic pTau, we show that vulnerability depends not only on upstream injury but on intrinsic neuronal capacity to contain it. L4 neuronal resilience to somatic tau accumulation was characterized by a greater proportion of pTau-resistant neuronal state and higher expression of resistance-associated programs, despite exposure to plaque-driven neuritic stress. Together with functional validation, these findings identify intrinsic neuronal state as a key determinant of Aβ-triggered neuronal pTau accumulation and shed light on the pathophysiological link between the two neuropathological hallmarks of AD. Targeting tau-vulnerability programs may therefore complement anti-amyloid therapies by limiting the progression from plaque-associated stress to somatic tau pathology before propagation occurs.

## ACKNOWLEDGEMENTS

We would like to thank the patients and families involved in research at the Massachusetts Alzheimer’s disease Research Center. The results published here are in whole or in part based on data obtained from the AD Knowledge Portal (https://adknowledgeportal.org). We acknowledge funding from the following sources: Alzheimer Association APOE Biology in Alzheimer’s (ABA) award ABA-25-1372179 to SD; Massachusetts Life Sciences Center Neuroscience Consortium; the Karen Toffler Charitable Trust, Cardinal Family Scholars Fund Cure Alzheimer’s Fund and NIH R01AG071567 (REB); NIH P30AG062421 to SD, ASP, and BTH.

## AUTHOR CONTRIBUTIONS

HL, REB, and SD conceived and designed the study. SD and HL wrote the initial draft of the manuscript. HL led all computational and statistical analyses. REB led the experimental studies, EG performed the functional validation assays, and A Devine performed the immunostaining assays. AG performed bioinformatics analyses of external datasets. SAP conducted the spatial transcriptomics assays. MWE, LG, and A Doering generated and analyzed the snRNA-seq data. TZ led the digital pathology analysis. BTH and AS-P contributed to data interpretation. All authors contributed to manuscript review and editing, and approved the final manuscript.

## METHODS

### Postmortem human brain tissue

Fresh-frozen postmortem human brain tissue from the inferior temporal gyrus (ITG; BA20) was obtained from the Massachusetts Alzheimer’s Disease Research Center (MADRC) under a protocol approved by the Mass General Brigham Institutional Review Board (IRB; protocol 1999P009556). Tissue samples from 12 donors, including Alzheimer’s disease donors stratified by APOE ε4/ε4 or ε3/ε3 genotype and control donors, were selected for this study. Detailed donor information is provided in Supplementary Table S1.

### Spatial transcriptomics using 10x Genomics Xenium In Situ

#### Initial quality control

Cryostat section (10 microns) was collected into a microcentrifuge tube and then RNA was extracted analyzed using an Agilent Tapestation. All tissues had RNA concentrations >250 pg/μl, DV200 >50%, and RIN >2. Adjacent cryostat sections were mounted on slides and Luxol Fast Blue Hemotoxylin & Eosin staining was performed to visualize tissue structure. Tissue sections were rehydrated by serial immersion in xylene, 100% ethanol, 95% ethanol, 70% ethanol, and distilled water (20 dips in each solution). Sections were then incubated in Luxol Fast Blue solution for 1 hr at 60°C, rinsed in distilled water, and counterstained with Harris Hematoxylin for 10 minutes at room temperature. Excess hematoxylin was removed with acid alcohol until nuclear staining was clearly visualized, followed by eosin incubation for 2 minutes at room temperature. Slides were then dehydrated through increasing concentrations of 70%, 95%, and 100% ethanol, followed by xylene (20 dips in each solution). Tissues were mounted with Eukitt Quick-hardening mounting medium and glass coverslips prior to being imaged on a VS120 slide scanner to assess the morphology and quality of staining. All tissues were free from freezing artefacts, with only small (cell-sized) holes in the tissue or were largely free from tissue disruption.

#### Xenium slide preparation and processing

Tissues that passed these quality control steps were sectioned from fresh frozen blocks onto Xenium slides according to the “Xenium In Situ - Fresh Frozen Tissue Preparation Handbook” (CG000579 Rev A, 10X Genomics) protocol. Tissue sections from four individual samples were distributed across two Xenium slides. The Xenium slides were then prepared following the “Xenium In Situ for Fresh Frozen Tissues - Fixation and Permeabilization” protocol (CG000581 Rev A, 10X Genomics). Briefly, the slides were heated at 37°C for 1 min and then successively submerged in 4% paraformaldehyde, PBS, SDS, and methanol to fix and permeabilize the tissue. Directly after, the samples were washed with PBS-T. The Xenium slides were then processed in accordance with the “Xenium Prime In Situ Gene Expression” user guide (CG000760 Rev A, 10X Genomics) for the rest of the slide preparation steps. The samples were incubated in priming hybridization mix with oligos from Xenium Prime 5K Human Pan Tissue & Pathways Panel (PN-1000671, 10X Genomics) and a custom-designed oligo set. Following priming hybridization, samples were treated with post-priming hybridization wash, RNase mix, and polishing mix, with washes between steps. The samples were then hybridized with probes from the Xenium Prime 5K Human Pan Tissue & Pathways Panel (PN-1000671, 10X Genomics) and a custom-designed probe set. Subsequently, samples were incubated in post-hybridization wash buffer, ligation mix, DNA amplification enhancement mix, and DNA amplification mix. The samples underwent cell segmentation staining and were then treated with autofluorescence quencher and DAPI before being loaded into the Xenium instrument. Processed Xenium slides were loaded into the Xenium Analyzer and imaged, following the guidelines in the “Xenium Analyzer User Guide (CG000584 Rev F, 10X Genomics).” After scanning, the Xenium slides were removed from the Xenium Analyzer. After all the *in situ* data was collected, sections were stored in phosphate buffered saline (PBS) at 4°C until additional immunofluorescent imaging was performed.

### Immunofluorescence Imaging

Tissues were briefly post-fixed with 4% paraformaldehyde (PFA) in PBS for 15 minutes and then were rinsed three times with PBS. Blocking solution (5% bovine serum albumin in PBS with 0.25% Triton-X) was applied for 30 minutes followed by incubation with primary antibodies in blocking solution overnight. These primaries included beta-amyloid (D54D2) rabbit monoclonal antibody (Alexa Fluor 488 conjugate; Cell Signaling Technology cat no. 51374; 1:500) and phospho-tau (Ser202, Thr205) mouse monoclonal antibody (AT8; custom Alexa Fluor 555 conjugate made from Invitrogen cat no. MN1020; 1:10,000). Sections were rinsed in PBS the next day and then coverslipped with Fluoromount G + DAPI (Southern Biotech) and imaged on an Olympus VS120 slide scanning microscope with a 20x objective lens. Following imaging, coverslips were gently removed in PBS, sections were rinsed and HS-84 (Sigma-Aldrich cat no. SML3751) was applied at 1:10,000 (from a 5 mg/ml stock) for one hour. Sections were rinsed extensively to remove excess dye in PBS, were coverslipped with Fluoromount G + DAPI and were re-imaged.

### Segmentation of histological features into masks

VS120 images were opened in ImageJ and the brightness was adjusted to normalize similar stained features having similar brightness range, then saved as a tiff with the adjusted pixel intensities. These adjusted images were then loaded into Ilastik^45^ and segmented following their pixel classifier and object classifier workflows. The pixel classifier workflow involves using a paintbrush to draw over signal and background to help train the classifier on how to segment each channel in each image. Pixel probability maps or segmentation maps were then exported. Pixel probability maps and raw data were then loaded into Ilastik’s object classification workflow, and segmented features were then classified as the object of interest or as noise. Data was then exported as a mask with all features from one class having equal pixel values (i.e. all pixels within a neurofibrillary tangle have a value of 1). Custom matlab code was then used to clean up the segmentation mask so each channel only has one classified object and all pixels not in that class are set equal to zero. Channels were then merged in ImageJ alongside a DAPI reference channel to allow alignment with spatial transcriptomics data.

### Xenium Spatial Transcriptomics raw data preprocessing and quality control

Raw Xenium outputs, including cell-by-gene count matrices, cell and nuclear segmentation boundaries, transcript coordinates and morphology/immunofluorescence images, were used for analysis. Data from individual samples were imported using spatialdata_io.xenium within the SpatialData^46^ framework (v0.3.0), retaining aligned images. Cell-by-gene count matrices were extracted as AnnData objects and concatenated across samples. Cells containing more than two unassigned codeword, negative-control codeword or negative-control probe transcripts were excluded to remove technical artifacts and background signal. Cells with an area outside 10 to 500 µm^2^, fewer than 15 detected transcripts or fewer than 10 detected genes were also excluded. Genes detected in fewer than 20 cells were removed from clustering analysis. In total, quality-control filtering excluded 2.15% of the cells.

We additionally evaluated transcript-informed segmentation using Baysor^47^ (v0.7.1) implemented through the Sopa framework (v2.0.0) using Xenium segmentation as prior. In this study, compared with the default Xenium segmentation, Baysor produced only marginal improvements in transcript inclusion and cell-identity assignment scores. Therefore, the default Xenium segmentation was used for downstream analyses.

### Major cell type and subtype annotations

Following quality control, the concatenated count matrix was processed using Scanpy^48^ (v1.11.0) in Python (v3.10.16). Counts were normalized to 1,000 transcripts per cell and log-transformed using log1p. The top 2,000 highly variable genes were identified using the Seurat method, scaled with values capped at 10, and subjected to principal component analysis. To correct for batch-associated variation, Harmony^49^ integration was applied to the principal component representation using *sample* identity as the batch variable. A nearest-neighbor graph was constructed using 30 neighbors and the first 20 Harmony-corrected principal components, followed by UMAP visualization and Louvain clustering at resolutions ranging from 0.1 to 0.5.

Major cell types were assigned using Harmony-corrected Louvain clusters at resolution 0.2, based on enrichment of brain cell-type marker genes. Marker sets were derived from a previously annotated SEA-AD middle temporal gyrus snRNA-seq reference dataset^50^ by differential expression analysis using rank_genes_groups, retaining genes using adjusted P < 0.01, log2 fold-change > 2, and expression in >60% of cells within the corresponding reference cell type. Log fold-change-weighted AUCell scores were calculated for each marker set using the aucell function in pySCENIC^51^ (v0.12.1), and clusters were annotated according to their predominant enrichment profiles. Cell type assignments were further validated by expression of canonical markers, including *MAP2* for neurons, *AQP4* for astrocytes, *CSF1R* for microglia, *CLDN5* for vascular cells, *MOG* for oligodendrocytes and *PDGFRA* for oligodendrocyte precursor cells (OPC).

Cell-subtype identities were assigned using Robust Cell Type Decomposition (RCTD) in spacexr^52^ (v2.2.1) in R (v4.4.1). Raw Xenium counts and cell centroid coordinates were used to construct spatial query objects. We used the cell subtype annotated snRNA-seq data^18,22,23^ from the same brain bank (MADRC) and cortical region (inferior temporal gyrus, ITG) as reference dataset. Batch-corrected cell-subtype mean expression signatures were estimated using scVI in cell2location^53^ (v0.1.3) with default parameters and supplied to RCTD as reference profiles. RCTD was run in doublet mode with a minimum UMI threshold of 0, a confidence threshold of 5 and a doublet threshold of 30. Assignments consistent with the corresponding major cell-type annotations were retained for downstream subtype-resolved spatial analyses.

### Cortical layer annotation

Cortical layers were annotated independently for each sample using the spatial organization of transcriptionally defined neuronal subtypes, followed by morphology-guided refinement using aligned DAPI, HuD and 18S rRNA images. Neurons with non-rejected RCTD assignments were used as anchors, with L2/3, L4, L5 and L6 identities defined by their excitatory neuronal subtype annotations. For each cell, spatial features were calculated as the mean Euclidean distance to the 20 nearest neurons overall and to the 20 nearest anchor neurons from each cortical layer. Putatively misplaced L6 anchors were excluded if their mean distance to L2/3 or L4 anchors was below the 98th percentile of the corresponding within-layer distance distribution.

Layer 1 and white matter were identified by integrating neuron depletion with k-means clustering (k = 8) based on scaled distances to L2/3, L4, L5 and L6 anchors. Neuron-depleted regions were defined as cells with a mean distance to the 20 nearest neurons exceeding the 95th percentile of that distribution among neuronal cells. Layers L2/3 to L6 were assigned using the RandomForestClassifier implementation in scikit-learn (v1.4.2), trained on the clean neuronal anchors, with distances to all neurons and to each layer-specific anchor population as features. Hyperparameters were selected by grid search with five-fold cross-validation, and the trained classifier was applied to all cells. Layer boundaries were manually refined in QuPath software (v0.5.1) based on tissue architecture and aligned DAPI, HuD and 18S rRNA staining patterns. Regions lacking coherent laminar organization or showing tissue artifacts were classified as noise and excluded from downstream analyses.

### Distance to plaque

The distance from cell boundary to the nearest plaque boundary was calculated using STRtree in Shapely (v2.0.6). Plaque objects were represented as polygons derived from the plaque mask and indexed with an STRtree. For each cell segmentation polygon, the nearest plaque polygon was queried, and the minimum two-dimensional Euclidean distance between the cell polygon and plaque polygon was calculated. Cells touching or overlapping a plaque were assigned a distance of zero.

### Definition of AT8^+^ pixels

AT8-positive pixels were identified using an adaptive intensity threshold anchored to a reference AD tissue section. AT8 pixel intensities within the tissue boundary of the reference section were transformed using log1p, and a robust upper-tail threshold was defined as the median plus 1.645 scaled median absolute deviations (MAD), corresponding to the upper 5% tail under a normal approximation:

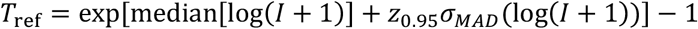

where *I* denote AT8 pixel intensity and *σ_MAD_* = 1.4826 × MAD is the MAD-based robust estimate of standard deviation.

To account for technical variation in overall fluorescence intensity across slides, the reference threshold was adjusted for each sample using the ratio of its mean tissue-level AT8 intensity to that of the reference sample, following the general adaptive-thresholding strategy previously described for immunofluorescence image analysis^54^:

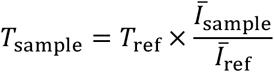

where *Ī*_sample_ and *Ī*_ref_ denote the mean within-tissue AT8 intensities in the corresponding sample and the reference sample, respectively. Pixels with AT8 intensity greater than or equal to *T*_sample_were classified as AT8^+^.

### Quantification of neuronal soma pTau burden

Soma pTau burden was quantified as an integrated measure of AT8 pathology within each neuronal soma boundary. For each soma, the summed AT8 intensity across AT8^+^ pixels was weighted by the fraction of the soma area occupied by AT8^+^ signal, and the resulting value was log-transformed:

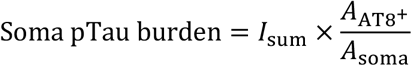

where *I_sum_* is the summed intensity across AT8^+^ pixels within the soma, *A*_AT8+_ is the AT8^+^ somatic area and *A*_soma_is the total soma area. Accordingly, this metric incorporates both the intensity and spatial extent of somatic pTau pathology while accounting for variation in soma size. Log-transformed values were used for downstream statistical analyses.

### Neurons pseudobulk differential gene expression

To identify differentially expressed genes (DEGs) associated with tau pathology-defined neuronal groups, we performed pseudobulk differential expression analyses for primary comparisons: (1) AD pTau-free versus Control neurons, and (2) AD Tangle+ versus AD pTau-free neurons. For each comparison, raw transcript counts were aggregated across individual cells within each donor, neuronal type, and defined comparison group, generating donor-by-neuron-type-by-group pseudobulk count matrices.

Normalization and differential expression testing were conducted using DESeq2^55^ (v1.44.0) in R (v4.4.1). For the unpaired comparison between pTau-free versus Control neurons, for each type of neurons, differential expression was assessed separately within each neuronal type using a negative binomial generalized linear model that adjusted for biological and technical covariates (design = ∼ Age + Sex + Batch + Comparison group). For within-donor comparisons (i.e., Tangle+ versus pTau-free neurons), we used a paired design to control for inter-donor variability (design = ∼ Donor + Comparison group). Effect sizes log2FoldChange were shrunken using apeglm^56^ method. P values were adjusted independently for each comparison using the Benjamini–Hochberg false-discovery rate (FDR) procedure. Genes meeting a significance threshold of FDR < 0.05 and abs(log2FoldChange) > 0.25 were considered significantly differentially expressed.

### Quantification of neuritic plaque-associated pTau burden

Neuritic plaque-associated (NP) pTau burden was quantified within plaque-centered regions of interest (ROIs), defined as the plaque core together with a 20 μm surrounding halo, after exclusion of all overlapping neuronal soma regions. For each ROI, the summed AT8 intensity across AT8^+^ pixels (described above), was weighted by the fraction of the ROI area occupied by AT8^+^ signal. The resulting integrated metric, hereafter referred to as NP pTau burden, was calculated as follows:

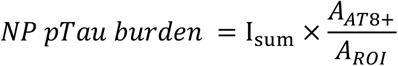

where *I*_sum_ is the summed AT8 intensity across AT8^+^ pixels, *A*_AT8+_is the AT8^+^ area and *A*_ROI_ is area of the plaque-centered ROI. Accordingly, this metric incorporates both AT8 signal intensity and the spatial extent of plaque-associated pTau pathology, while accounting for ROI area.

To assess whether this metric primarily reflected local pTau pathology rather than plaque size, principal component analysis was performed using plaque size, mean AT8 intensity within AT8^+^ pixels and the AT8^+^ area fraction of the plaque ROI as input features.

### Identification of genes associated with NP pTau burden in neuropil and glia

Neuritic plaque-associated pTau (NP pTau) burden was modeled as a continuous variable and standardized within donor before analysis. For the neuropil analysis, transcript counts were obtained from the 20-µm plaque halo after exclusion of regions overlapping with cell somata. For the glial analyses, astrocytes and microglia located within the corresponding 20-µm plaque halo were analyzed separately. Each neuropil region of interest (ROI) or glial cell was assigned the NP pTau burden and cortical layer annotation of its associated plaque.

Data processing was performed in Python (v3.10.16) using Scanpy (v1.11.0). For each analysis, count matrices were normalized to 1,000 transcripts per neuropil ROI or glial cell and log-transformed using log1p. Genes detected in at least 10% of neuropil ROIs or cells in the corresponding analysis were retained. For each retained gene, normalized expression was modeled using ordinary least squares as implemented using statsmodels.formula.api.ols in statsmodels (v0.14.4): Gene expression ∼ NP pTau burden (scaled) + cortical layer + donor.

For the glial-cell analyses, plaque-clustered robust standard errors were used because multiple cells could be associated with the same plaque and therefore share the same NP pTau burden measurement. Donor was included as a fixed effect considering the small number of donors analyzed (n = 4) does not support reliable estimation of a donor-level random-effect variance. P values were adjusted separately for the neuropil, astrocyte and microglial analyses using the Benjamini–Hochberg false-discovery rate (FDR) procedure.

### SEM modelling NP pTau burden effect on neuritic stress

Plaque-level structural equation modelling (SEM) was performed using plaques in cortical layers L2/3, L4, L5 and L6 with available NP pTau burden, neuropil neuronal stress score, microglial response score and astrocytic response score.

Microglial and astrocytic response signatures were derived from the corresponding NP pTau association analyses described above. For each cell type, genes expressed in more than 10% of cells, with an absolute regression coefficient > 0.01 and nominal P < 0.05, were separated into positively and negatively associated gene sets. Marker genes of other cell types were excluded. Module scores for these gene sets were calculated in microglia or astrocytes within the 20 μm plaque halo using AddModuleScore in Seurat (v5.1.0). For each plaque, the microglial response score was calculated as the mean positive-module score minus the mean negative-module score across halo-localized microglia. The astrocytic response score was calculated analogously. The neuropil neuronal stress signature was defined using neuron-enriched genes positively associated with NP pTau burden in the neuropil analysis described above, after exclusion of non-neuronal contaminant genes. A neuropil neuronal stress score was calculated for each plaque halo using AddModuleScore in Seurat^57^.

NP pTau burden, neuropil neuronal stress score, microglial response score and astrocytic response score were standardized within donor before modelling. SEM was performed in R using lavaan^58^ (v0.6.21), with NP pTau burden as the exposure, microglial and astrocytic response scores as parallel mediators, and neuropil neuronal stress score as the outcome. The model included both direct effect from exposure to outcome and indirect effects mediated through microglial or astrocytic. Residual covariance between the two mediator variables was included in the model. The proportion of the total effect represented by each direct or indirect pathway was calculated by dividing that effect by the total effect. Standardized effect estimates, P values and 95% confidence intervals were obtained using 2,000 bootstrap iterations.

### Distance-decay modeling of neuron-level plaque toxicity exposure

We quantified each neuron’s NP pTau exposure, referred to as plaque toxicity exposure, as the distance-weighted sum of NP pTau burden from all plaques within 100 μm of the cell, using an exponential distance-decay model.

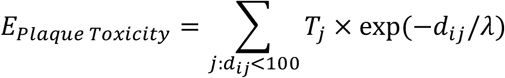

where *T_j_* denotes the NP pTau burden of plaque j, *d_ij_* denotes the Euclidean distance between neuron i and plaque j, and λ denotes the distance-decay parameter, in this study, λ was set to 20 μm.

### Association between NP pTau exposure and soma pTau

We restricted the analysis to AD excitatory neurons in cortical layers L2/3, L4, L5 and L6 that had at least one plaque within 100 μm. For each neuron, plaque toxicity exposure was log-transformed and standardized within donor. Its association with somatic pTau burden was tested using a linear mixed-effects model:

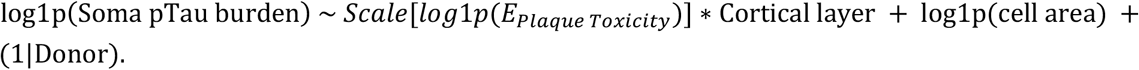

This model estimated the relationship between plaque toxicity exposure and somatic tau burden while adjusting for cell size and donor-level variation and allowed the exposure slope to differ across cortical layers. Layer-specific associations between plaque toxicity exposure and somatic pTau burden were estimated from the fitted model using emtrends in emmeans (v2.0.2), together with its standard error, confidence interval and P value.

Given the limited number of donors, which may lead to unstable estimation of random-effect variance, we additionally performed a sensitivity analysis modeling donor identity as a fixed effect. This analysis yielded nearly identical results.

### Neuronal trajectory and branch-associated gene expression analysis

Neuronal trajectory analyses were performed separately for L2/3, L4 and L5 excitatory neurons from AD Xenium spatial transcriptomic samples. Trajectories were constructed using gene expression profiles only, and pathology measurements were mapped onto the inferred trajectories only after trajectory construction. For each type of neurons, genes detected in fewer than 50 cells in the spatial transcriptomic dataset, expressed in less than 5% of the corresponding neuronal population in the same region snRNA-seq reference dataset, or predefined as non-neuronal marker genes were excluded, thereby restricting the analysis to detected neuronal transcripts.

To prepare input for trajectory construction, transcript counts were normalized to 1,000 transcripts per cell and log-transformed using Scanpy (v1.11.0). Highly variable genes (HVGs) were identified from the raw count layer using the Seurat v3 method implemented in Scanpy, with donor identity specified as the batch variable. The top 1,000 HVGs were considered, and only genes identified as highly variable in at least three donors were retained. Log-normalized expression values for the retained HVGs were scaled with values capped at 10, followed by principal component analysis (PCA). To minimize donor-associated variation while preserving within-layer neuronal heterogeneity, Harmony integration was applied to the PCA embedding using donor identity as the integration variable.

Trajectory inference was performed using Monocle 2^59^ (v2.32.0). The first 20 Harmony-corrected principal components were used as low-dimensional input for DDRTree reconstruction, rather than raw gene-expression values, to define trajectories from donor-corrected neuronal-state variation. A Monocle CellDataSet was constructed using a Gaussian-family model, and no additional normalization or scaling was applied during DDRTree embedding. DDRTree was run with two output dimensions and 100 centroids (for L4 excitatory neurons, we used 30 centroids). Cells were ordered along the resulting trajectory, and pseudotime was rooted at the selected starting state. The branch point of interest was selected from the reconstructed trajectory for downstream analysis. Each neuron was assigned its corresponding somatic pTau burden and neuritic plaque-associated pTau exposure value, quantified as described above.

To identify genes with branch-associated expression dynamics, branch assignments and pseudotime inferred from the Harmony-corrected trajectory were transferred to an expression-based Monocle CellDataSet containing the corresponding neuronal transcript-count data. To reduce the disproportionate contribution of densely represented terminal states, cells were stratified into 50 pseudotime bins and randomly downsampled to a maximum of 100 cells per bin using a random seed of 42. Genes detected in at least 200 retained cells were tested using branched expression analysis modeling (BEAM) function implemented in Monocle. For each gene, the full model was specified as expression ∼ sm.ns(Pseudotime, df = 3) × Branch + Donor, and the reduced model was specified as expression ∼ sm.ns(Pseudotime, df = 3) + Donor. Genes with a BEAM false-discovery rate-adjusted q value < 0.05 and successful model fitting status were considered to exhibit significant branch-associated expression dynamics.

To compare transcriptional states between trajectory branches, we performed pseudobulk differential expression analysis separately for each neuronal layer. Raw Xenium transcript counts were aggregated by donor and branch state to generate donor-level pseudobulk profiles. Differential expression analysis was performed using DESeq2 (v1.44.0), with donor identity included in the design formula to account for paired inter-donor variation (design = ∼ Donor + State). Genes meeting a significance threshold of FDR < 0.05 and abs(log2FoldChange) > 0.25 were considered significantly differentially expressed.

### Functional enrichment analysis

Functional enrichment analyses were performed in R (v4.4.1). For gene set enrichment analysis (GSEA), genes from each differential expression comparison were ranked by the corresponding test statistic. Gene sets for Gene Ontology Biological Process (GOBP), Cellular Component (GOCC) and Reactome pathways were obtained from the Molecular Signatures Database (MSigDB) using msigdbr^60^ (v24.1.0). GSEA was performed using the GSEA function in clusterProfiler^61^ (v4.12.6) with a minimum gene set size of five genes and a fixed random seed of 123. Enrichment results were assessed using Benjamini–Hochberg false-discovery rate (FDR)-adjusted P values.

For over-representation analysis, significantly differentially expressed genes were converted from gene symbols to Entrez identifiers using bitr and org.Hs.eg.db (v3.19.1). GOBP enrichment was performed using enrichGO in clusterProfiler package (v4.12.6), and Reactome pathway enrichment was performed using enrichPathway in ReactomePA^62^ (v1.48.0). Analyses were conducted separately for upregulated and downregulated genes, using all genes tested in the corresponding differential expression analysis as the background gene universe.

## DATA AVAILABILITY

The spatial transcriptomics dataset generated for the study will be made available at Zenodo (https://doi.org/10.5281/zenodo.21134530; https://doi.org/10.5281/zenodo.21042158; https://doi.org/10.5281/zenodo.21134777). Genes and pathways information are available as Supplementary Data.

snRNA-seq data used in this study can be found in the Synapse, GEO, or custom public repositories as follows: AD progression study which has matched snRNA-seq from same donors (Zenodo: 13222113; GEO: GSE268599; SRA: PRJNA1142727, PRJNA916657;), SEA-AD (MTG region, processed nuclei h5ad file: https://sea-ad-single-cell-profiling.s3.amazonaws.com/MTG/RNAseq/Reference_MTG_RNAseq_final-nuclei.2022-06-07.h5ad), Mathys study ROSMAP (Synapse: syn52293417), Gazestani brain-biopsy cohort (available at: https://braincelldata.org/resource).

## CODE AVAILABILITY

All code generated during this study is accessible at https://github.com/mindds/AD-spatial-tau-vulnerability.

## SUPPLEMENTARY TABLES

1. TableS1_Donor_Characteristics.xlsx

2. TableS2_AD-pTau-free_vs_Ctrl.pbulk.xlsx

3. TableS3_NP_pTau_assoc_InHalo-Neuropil_genes.xlsx

4. TableS4_NP_pTau_assoc_InHalo-Astrocytes_soma_genes.xlsx

5. TableS5_NP_pTau_assoc_InHalo-Microglia_soma_genes.xlsx

6. TableS6_Soma_pTau_assoc_L2-3-L5_Ex_soma_genes.xlsx

7. TableS7_Trajectory_State2_vs_state3.DESeq2_across_layer_ex.xlsx

8. TableS8_APOE_e4_vs_e3_pathway_analysis.xlsx

---

**Extended Data Fig 1.**
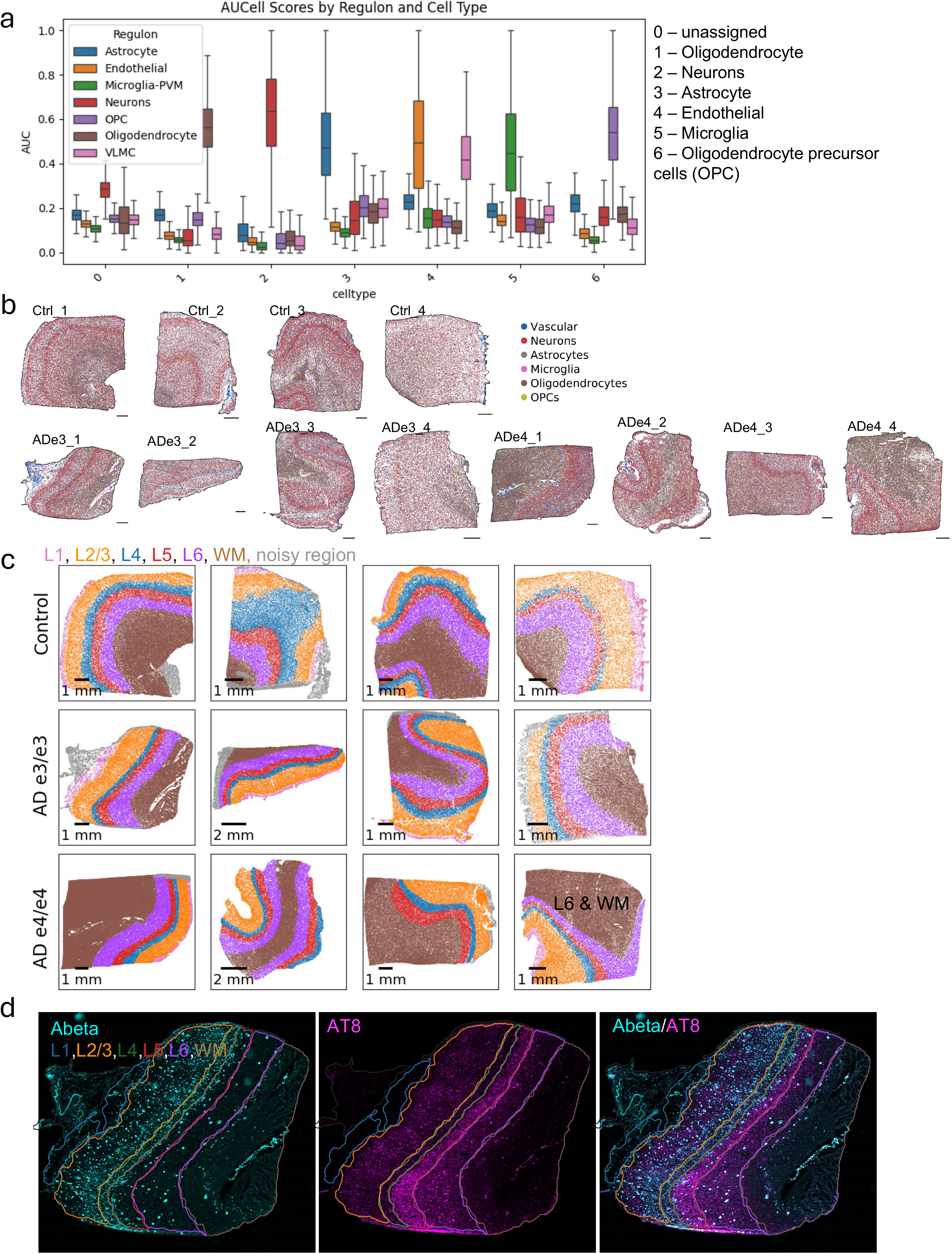
**a**, Cell types were annotated by enrichment of brain cell-type marker sets derived from the SEA-AD dataset. Cluster 1 was assigned to oligodendrocytes, cluster 2 to neurons, cluster 3 to astrocytes, cluster 4 to vascular-associated cells, cluster 5 to microglia and cluster 6 to oligodendrocyte precursor cells (OPCs). **b**, Spatial maps showing cell-type annotations across control (Ctrl), AD *APOE* ε3/ε3 (ADe3), and AD *APOE* ε4/ε4 (ADe4) samples. **c**, Spatial maps showing cortical layer annotations from L1 to L6 and white matter (WM) in Ctrl (first row) and *APOE* ε3/ε3 (second row) and AD *APOE* ε4/ε4 (third row) samples. **d**, Representative Aβ and AT8 immunofluorescence images of Alzheimer’s disease inferior temporal gyrus (ITG) tissue, with cortical layers L1–L6 and white matter (WM) annotated.

**Extended Data Fig 2.**
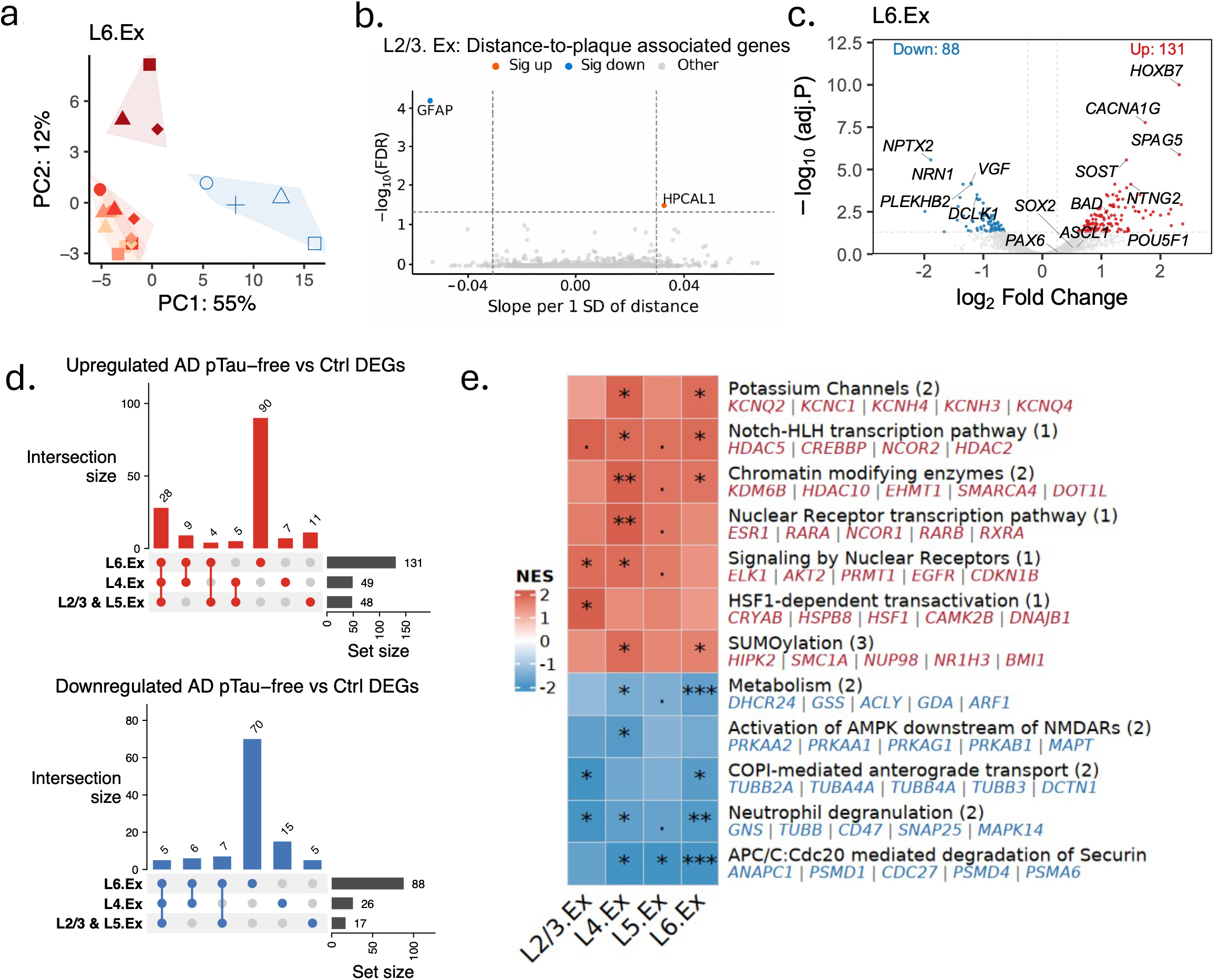
**a**, PCA of L6 excitatory neuron pseudobulk transcriptomes, related to Figure 2a. **b**, Volcano plot showing distance-to-plaque-associated genes in L2/3 excitatory neuron somata. Sig up: FDR < 0.05, slope ≥ 0.03; Sig down: FDR < 0.05, slope ≤ -0.03. **c**, Volcano plots showing differential gene expression between AD pTau⁻ and CTRL excitatory neurons in L6. **d**, UpSet plots showing overlap of upregulated and downregulated genes in AD pTau-free versus control excitatory neurons across cortical layers. **e**, Heatmap of Reactome pathway enrichment in AD pTau⁻ vs. CTRL excitatory neurons across cortical layers. Redundant pathways were clustered by leading-edge gene overlap (overlap coefficient > 0.7), with the most significant pathway representing each cluster. Colors indicate normalized enrichment scores (NES), and asterisks indicate FDR significance (\*\*\**q* < 0.001, \*\**q* < 0.01, \**q* < 0.05, ^•^*q* < 0.1).

**Extended Data Fig 3.**
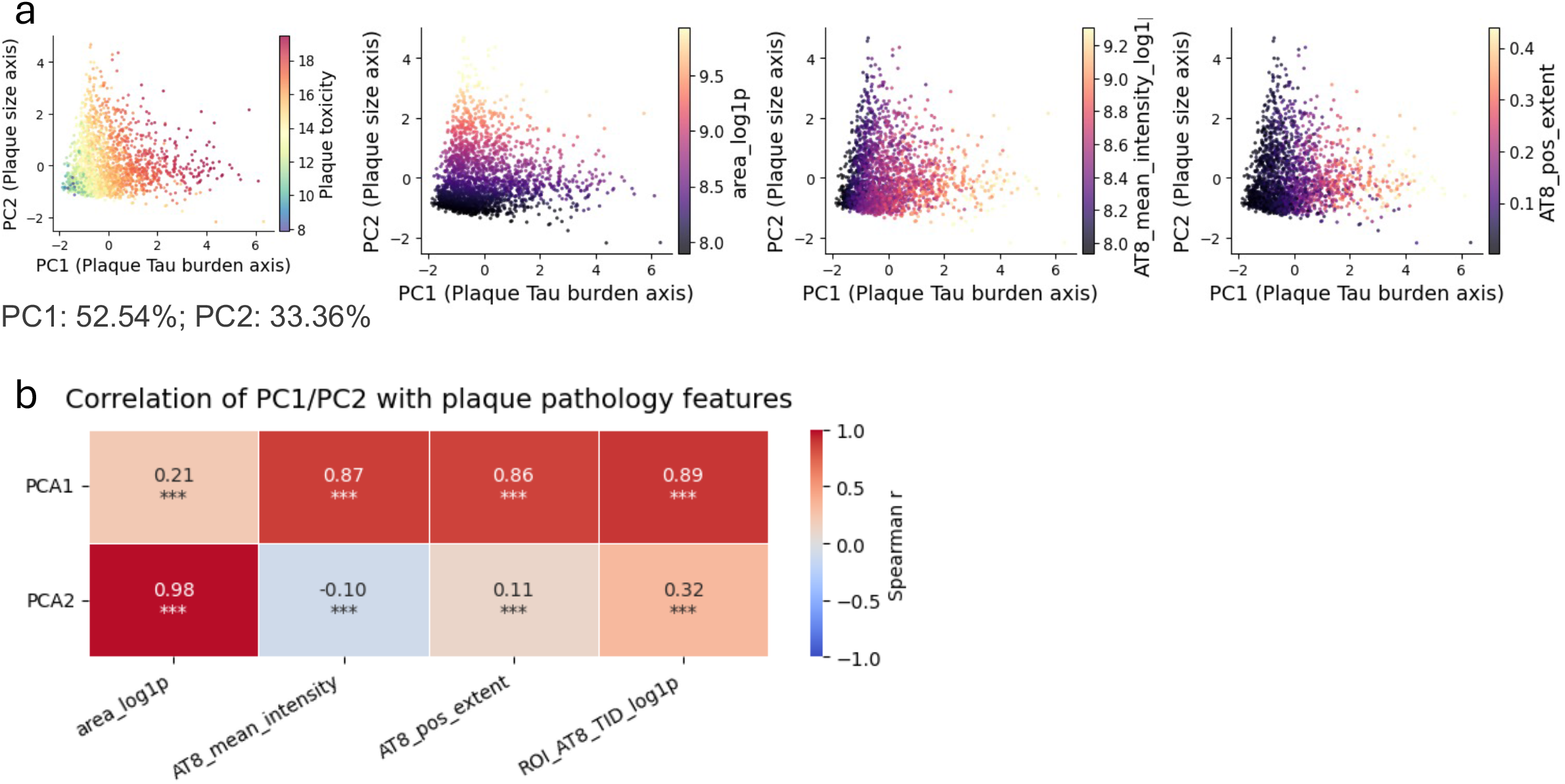
**a**, Principal component analysis of plaque-level pathology features measured within plaques and their surrounding 20 μm halos. Each point represents an individual plaque. Points are colored by calculated NP pTau, plaque area, mean AT8 intensity and AT8-positive extent. **b,** Spearman correlation heatmap showing associations between PC1 or PC2 and plaque pathology features, including plaque area, mean AT8 intensity, AT8-positive extent and integrated neuritic-plaque pTau burden (calculated NP pTau). PC1 was strongly correlated with AT8 intensity, AT8-positive extent and integrated neuritic-plaque pTau burden, and only weakly correlated with plaque area. ***P < 0.001.

**Extended Data Fig 4.**
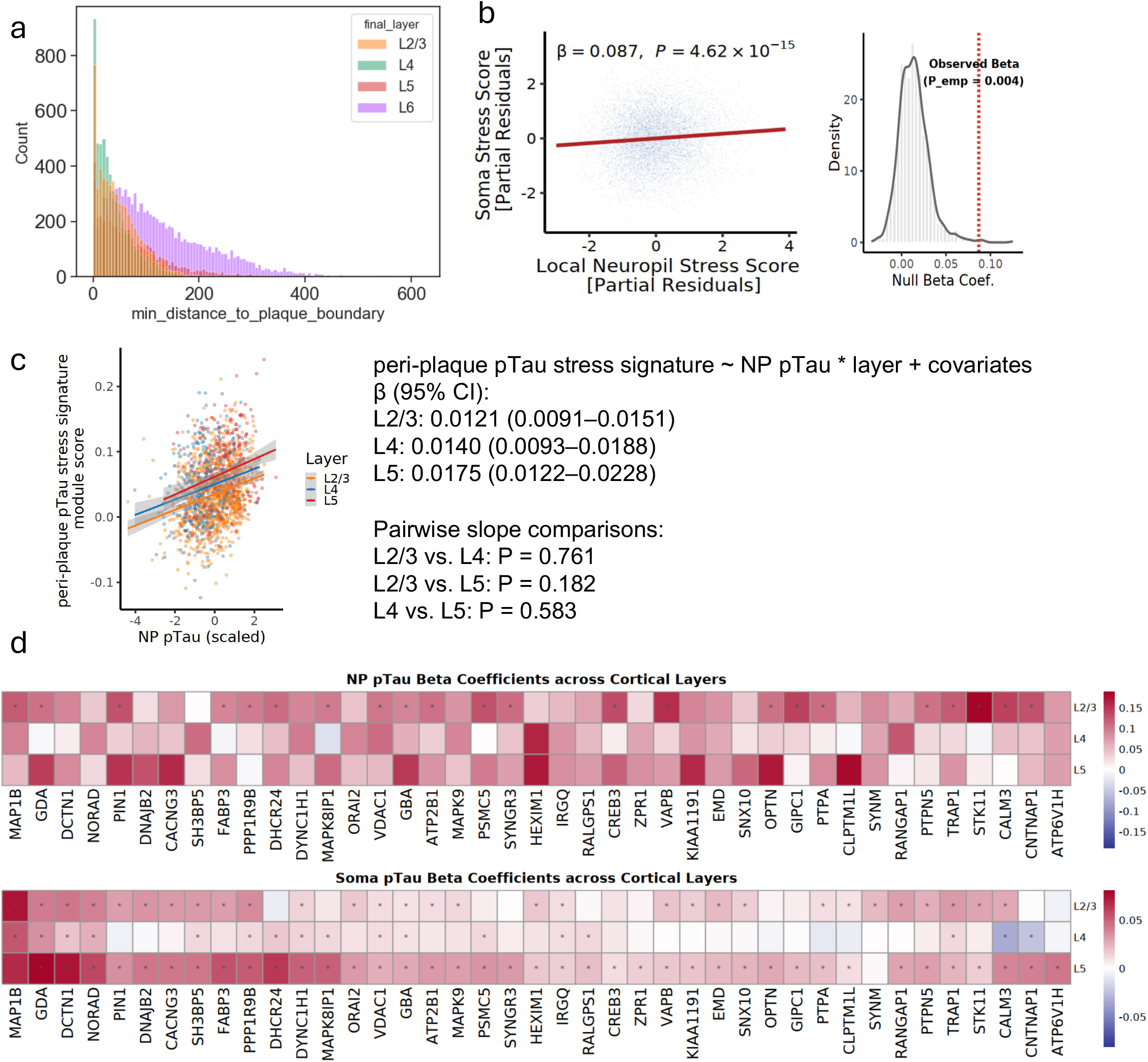
**a**, Distribution of the minimum distance from each excitatory neuron soma to the nearest plaque boundary, stratified by cortical layer. **b,** (left) Association between the local neuropil stress score and the somatic stress score across L2/3 and L5 excitatory neurons after covariate adjustment. (right) The corresponding permutation-based null distribution of regression coefficients, generated from 1,000 random gene sets; the observed coefficient is indicated by the red dashed line. **c,** Lines show layer-specific linear mixed-effects model fits adjusted for plaque area, with donor included as a random intercept. Shaded regions indicate 95% confidence intervals. Slopes did not differ significantly between layers (all pairwise P > 0.18). **d**, Layer-stratified regression coefficients for genes in the shared neurite–soma stress program. Heatmaps show associations between NP pTau and gene expression in the local neuropil compartment (left) or somatic pTau burden and gene expression in the soma compartment (right), across L2/3, L4 and L5 excitatory neurons. Colors indicate regression beta coefficients. Asterisks indicate significant associations. *p < 0.05.

**Extended Data Fig 5.**
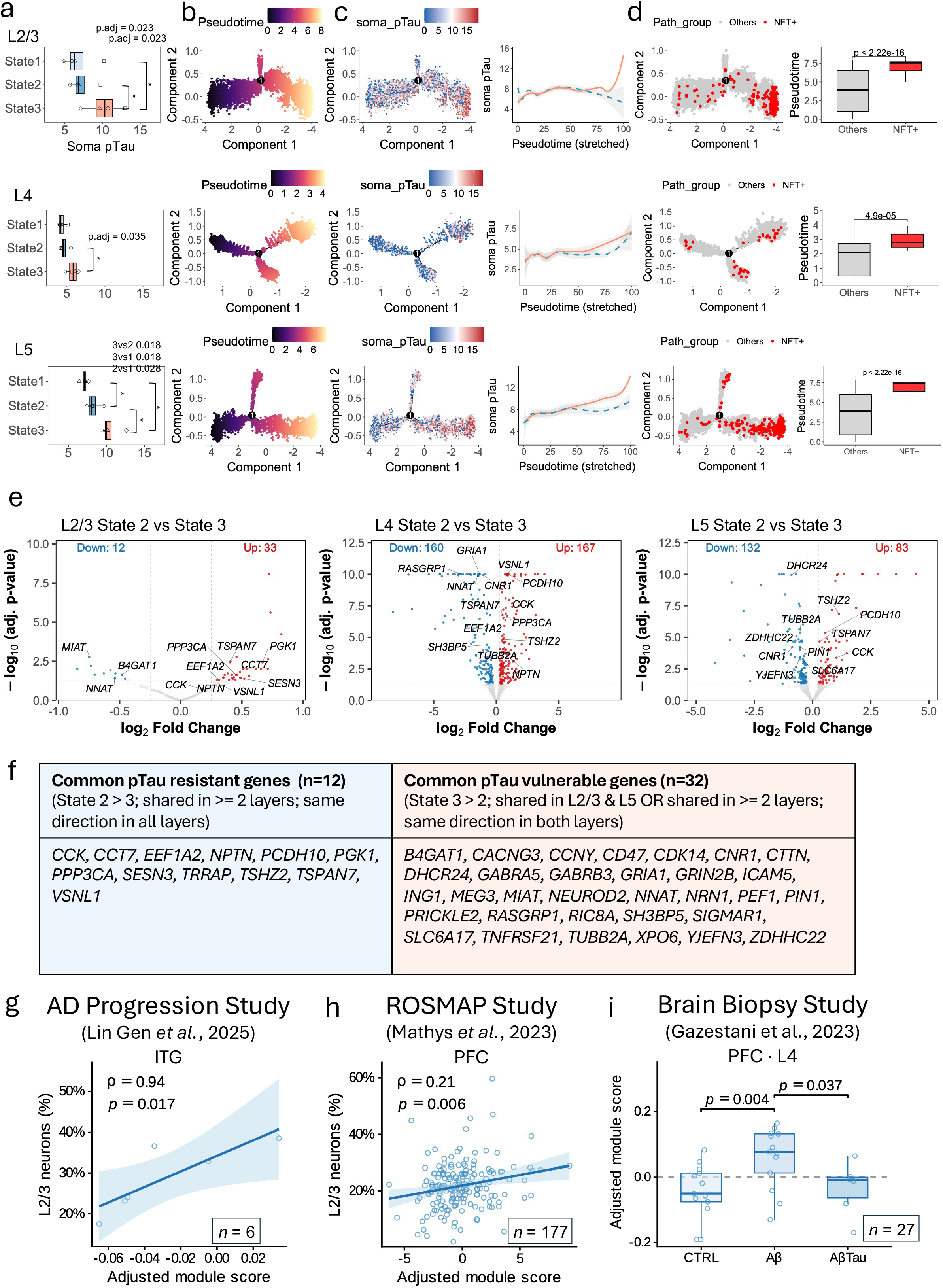
**a,** Plot showing donor-level somatic pTau burden across trajectory-defined states in L2/3, L4 and L5 excitatory neurons. Each point represents one donor, with point shape denoting donor identity. Adjusted *P* values for pairwise comparisons are indicated. **b,** Monocle trajectories inferred from transcriptomic profiles alone for L2/3, L4 and L5 excitatory neurons, colored by pseudotime. Black circles mark the branch point/root state. **c,** Same trajectories colored by somatic pTau burden (left), with branch-resolved trends in soma pTau plotted along stretched pseudotime (right). Solid lines indicate the State 1-to-State 3 trajectory, whereas dashed lines indicate the State 1-to-State 2 trajectory. **d,** Distribution of tangle-bearing neurons (NFT+) along each trajectory (left) and comparison of pseudotime between NFT+ and all other neurons (right). NFT+ neurons localized to the distal end of the pTau-vulnerable branch and occupied significantly later pseudotime than non-tangle-bearing neurons. **e,** Volcano plots showing differential expression between the pTau-resistant State 2 and pTau-vulnerable State 3 within L2/3, L4 and L5 excitatory neurons. Representative differentially expressed genes are labeled. Numbers indicate the total genes significantly decreased (blue) or increased (red) in State 2 relative to State 3. **f,** Cross-layer summary of genes associated with resistant and vulnerable states. Left, gene symbols enriched in the common pTau-resistant program (State 2 > State 3) and genes enriched in pTau-vulnerable states (State 3 > State 2). Right, functional grouping of representative common pTau-resistant genes (blue) and pTau-vulnerable genes (orange). **g,** AD Progression Study: Association between the L2/3 excitatory pTau-resistant module score, adjusted for age at death, sex, and PMI, and the observed proportion of L2/3 neurons among all neurons in ITG donors with Braak stages II–III and sparse or moderate CERAD neuritic plaques (n = 6). Points represent donors; the line and shading represent the linear regression fit and 95% confidence band. **h,** ROSMAP Study: Association between the L2/3 excitatory pTau-resistant module PC1 score, adjusted for age at death, sex, PMI, education, and APOE ε2 and ε4 carrier status, and the observed proportion of L2/3 neurons among all neurons in PFC donors with Braak stages III–IV (n = 177). Points represent donors; the line and shading represent the linear regression fit and 95% confidence band. **i,** Brain Biopsy Study: Age- and sex-adjusted scores for the common pTau-resistant gene module in L4 RORB_SCTR+ neurons across control (CTRL), amyloid-positive (Aβ), and amyloid- and tau-positive (AβTau) donors (n = 27). Boxes show the median and interquartile range, points represent donors, and brackets indicate nominal pairwise *p*-values from estimated marginal mean contrasts following ANCOVA adjusted for age and sex. **Abbreviations:** ANCOVA, analysis of covariance; ITG, inferior temporal gyrus; PC1, first principal component; PFC, prefrontal cortex; PMI, postmortem interval

